# RHINO restricts MMEJ activity to mitosis

**DOI:** 10.1101/2023.03.16.532763

**Authors:** Alessandra Brambati, Olivia Sacco, Sarina Porcella, Joshua Heyza, Mike Kareh, Jens C. Schmidt, Agnel Sfeir

## Abstract

DNA double-strand breaks (DSBs) are toxic lesions that can lead to genome instability if not properly repaired. Breaks incurred in G1 phase of the cell cycle are predominantly fixed by non-homologous end-joining (NHEJ), while homologous recombination (HR) is the primary repair pathway in S and G2. Microhomology-mediated end-joining (MMEJ) is intrinsically error-prone and considered a backup DSB repair pathway that becomes essential when HR and NHEJ are compromised. In this study, we uncover MMEJ as the major DSB repair pathway in M phase. Using CRISPR/Cas9-based synthetic lethal screens, we identify subunits of the 9-1-1 complex (RAD9A-HUS1-RAD1) and its interacting partner, RHINO, as critical MMEJ factors. Mechanistically, we show that the function of 9-1-1 and RHINO in MMEJ is inconsistent with their well-established role in ATR signaling. Instead, RHINO plays an unexpected and essential role in directing mutagenic repair to M phase by directly binding to Polymerase theta (Polθ) and promoting its recruitment to DSBs in mitosis. In addition, we provide evidence that mitotic MMEJ repairs persistent DNA damage that originates in S phase but is not repaired by HR. The latter findings could explain the synthetic lethal relationship between *POLQ* and *BRCA1/2* and the synergistic effect of Polθ and PARP inhibitors. In summary, our study identifies MMEJ as the primary pathway for repairing DSBs during mitosis and highlights an unanticipated role for RHINO in directing mutagenic repair to M phase.

---

MMEJ is an intrinsically mutagenic repair pathway. Nevertheless, it mitigates the harmful effects of double-strand breaks (DSBs) by preventing the accumulation of large-scale DNA rearrangements. Furthermore, repair by MMEJ is necessary for the survival of cells with compromised HR and NHEJ (Ceccaldi *et al*, 2015; Mateos-Gomez *et al*, 2015; Wyatt *et al*, 2016). As a result, targeting this pathway has emerged as a promising therapeutic approach for the increasing number of cancer patients with defective HR, including ones carrying mutations in *BRCA1* and *BRCA2* (Schrempf *et al*, 2021; Zatreanu *et al*, 2021; Zhou *et al*, 2021). MMEJ is characterized by the presence of 2-6 base pairs (bp) of microhomology, as well as insertions and deletions (indels) that scar repair sites (Sfeir & Symington, 2015). These indels are introduced by DNA polymerase theta (Polθ), encoded by the *POLQ* gene. Polθ is a low-fidelity polymerase with a helicase-like activity that plays a central role in MMEJ (Audebert *et al*, 2004; Black *et al*, 2019).

The mutational signature associated with MMEJ has been found across different species, and the pathway is conserved from bacteria to humans (Ramsden *et al*, 2022). However, its mechanistic basis remains poorly defined. In mammalian cells, studies have demonstrated that following the formation of a DSB, short-range DNA end-resection by MRE11 and CtIP exposes flanking microhomologies that promote the annealing of opposite ends of the break (Lee-Theilen *et al*, 2011; Truong *et al*, 2013; Xie *et al*, 2009). When internal homologies are base-paired, they form single-stranded DNA (ssDNA) flaps that are cleaved by APEX2 and FEN1 (Fleury *et al*, 2022; Mengwasser *et al*, 2019; Sharma *et al*, 2015). Annealed intermediates are extended by Polθ (Arana *et al*, 2008; Chan *et al*, 2010) and sealed by XRCC1/LIG3 to complete end-joining (Audebert *et al*., 2004). Polθ also acts on transient “snap-back” substrates formed when the overhang of resected DSBs folds back and anneals to itself. Ultimately, both Polθ-mediated insertions contribute to the mutagenicity of MMEJ (Kent *et al*, 2016). While upregulated in many cancer types, Polθ is generally low in abundance and must be actively recruited to DSB sites (Seki *et al*, 2003). Nevertheless, how Polθ gets targeted to break sites and the upstream factors that drive MMEJ remain unknown.

MMEJ was initially discovered as an inefficient joining activity in Ku-deficient *Saccharomyces cerevisiae* (Boulton & Jackson, 1996). In higher eukaryotes, MMEJ is essential only when HR and NHEJ are blocked. Therefore, it emerged as a backup pathway that acts in the absence of the preferred modes of DSB repair (Ceccaldi *et al*., 2015; Mateos-Gomez *et al*., 2015; Wyatt *et al*., 2016). However, recent reports suggest that under certain conditions, MMEJ prevails. For instance, MMEJ is the primary repair mechanism for CRISPR/Cas9-induced DSBs in early zebrafish embryos (Thyme & Schier, 2016). In human and mouse cells, studies have reported that MMEJ occurs with NHEJ to promote random integration of foreign DNA into the genome and when repairing CRISPR/Cas9-induced breaks in particular loci (Corneo *et al*, 2007; Saito *et al*, 2017; Schimmel *et al*, 2017). Despite these findings, the dedicated physiological function of MMEJ and when cells opt for mutagenic MMEJ repair is poorly understood.

Polθ inhibitors are currently in phase I/II in the clinic as monotherapy and in conjunction with PARP inhibitors (PARPi) (ClinicalTrials.gov Identifier: NCT04991480). Preclinical studies demonstrated that Polθ inhibitors target BRCA-defective tumors and display significant synergy with PARPi and can effectively eliminate some PARPi-resistant tumors (Monica Bubenik, 2022; Zatreanu *et al*., 2021; Zhou *et al*., 2021). Elucidating the underlying mechanism of MMEJ and its temporal and spatial regulation is critical to understanding when and how cells choose to use mutagenic MMEJ and potentially explain the synthetic lethal interaction between MMEJ and HR.

## Results

### CRISPR/Cas9 synthetic lethal screen uncovers the full spectrum of MMEJ factors

To identify potential MMEJ factors, we conducted a genome-wide CRISPR-based synthetic lethal screen in cells that lack both HR and NHEJ. We hypothesized that since cells lacking these canonical DSB repair pathways are highly dependent on MMEJ for survival, this approach would reveal the full spectrum of MMEJ factors (Fig. 1a). We used CRISPR/Cas9 targeting in *BRCA2*^-/-^ DLD1 cells and generated clonally derived BRCA2^-/-^LIG4^-/-^TP53^-/-^ cells (referred to as TKO for triple knockout cells; Fig. 1b and Fig. S1A-B). To functionally validate the TKO cells, we treated them with a small molecule inhibitor of Polθ, RP6685 (Monica Bubenik*, 2022), and observed the expected loss in viability (Fig S1C). We then transduced TKO and TP53^-/-^ cells with the Toronto Knock Out version3 library targeting approximately 18,000 genes with four guides per gene (Hart *et al*, 2017). Cells were collected shortly after recovery from selection and after 14 population doublings. The integrated sgRNA were then subjected to targeted next-generation sequencing (NGS), and the relative sgRNA abundance was computed using the BAGEL2 software to yield an essentiality Bayes Factor (BF) score for each gene (Fig. 1c) (Kim & Hart, 2021). Common essential genes were depleted similarly in TKO and TP53^-/-^ cells, thus confirming the screen’s performance (Fig. S1D).

**Figure. 1.**
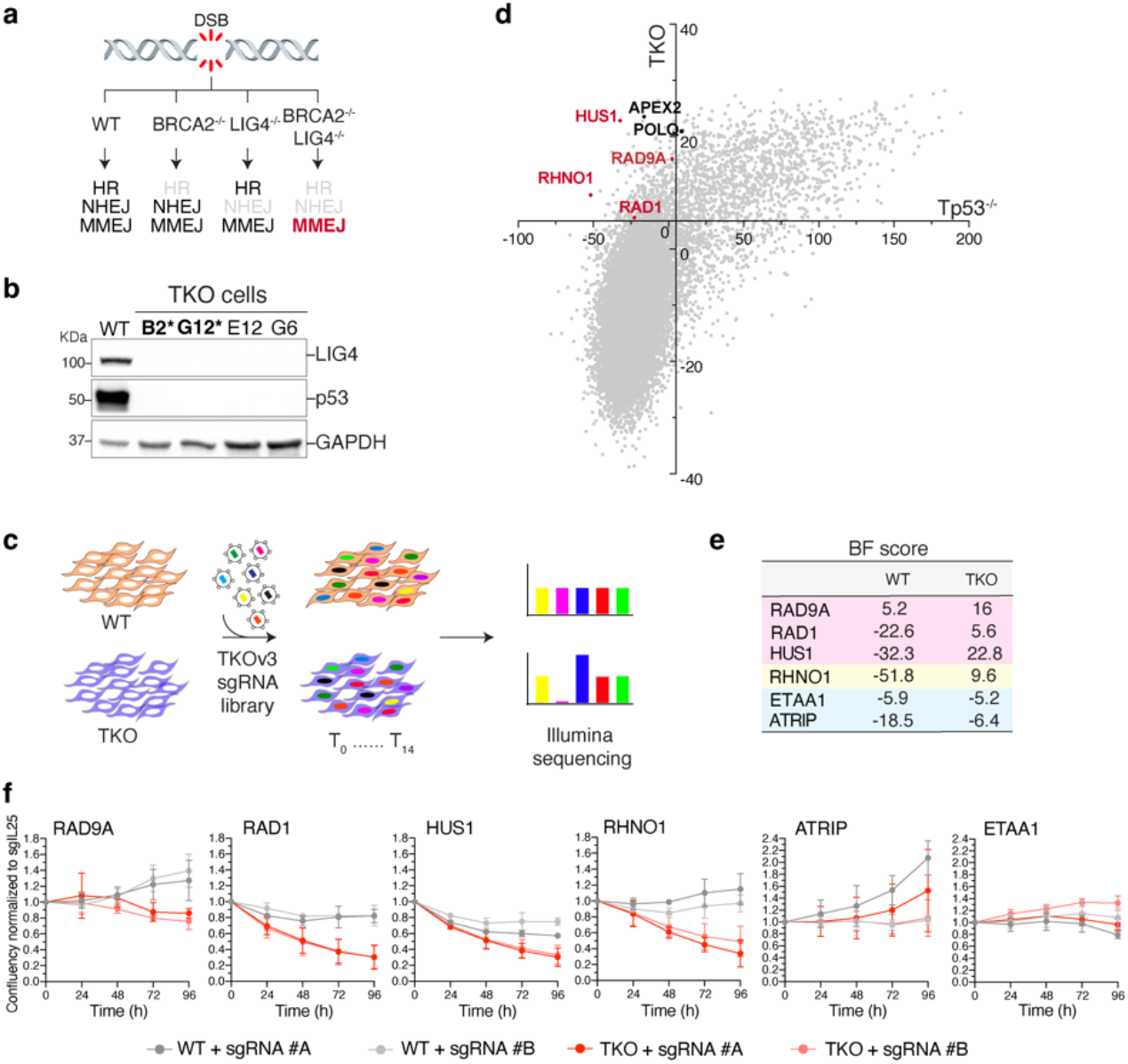
A genome-wide CRISPR-Cas9 screen uncovers an essential function for 9-1-1 and RHNO1 in cells lacking BRCA2 and LIG4. Schematic representation of the three major DSB repair pathways in mammalian cells: MMEJ, NHEJ, and HR. (**B**) Western blot analysis of LIG4 and p53 in clonally derived BRCA2^-/-^ LIG4^-/-^ TP53^-/-^ DLD1 cells (TKO). The asterisk indicates the clones used in the screen. (**C**) Schematic of the dropout CRISPR/Cas9 screen to identify synthetic lethal interactions. (**D**) Result of the genome-wide CRISPR-Cas9 screen in TKO cells compared to control cells. Bayes factor (BF) scores for essential genes in TKO vs. WT cells. BF scores were determined using the BAGEL2 analysis pipeline. Highlighted in red are subunits of the 9-1-1 complex, RHNO1, and in black MMEJ factors. The screen dataset is available in Supplementary Table 1. (**E**) Table reporting the BF score for the indicated genes. (**F**) Growth curve of TKO and WT cells treated with the indicated sgRNAs. Data are mean ± s.d. of three independent experiments normalized first to the initial time point (one day after seeding) and then to control sgRNA (sgIL25).

Our genome-wide screen identified a set of genes that were preferentially depleted in TKO cells, including hits previously reported to be essential for the survival of *BRCA2* null cells, such as *FEN1, RNaseH2A/B/C* (Zimmermann *et al*, 2018), *CIP2A* (Adam *et al*, 2021), and *ALC1* (Verma *et al*, 2021) (Fig. S1E), and known MMEJ factors, including *POLQ, HMCES* (Shukla *et al*, 2020), and *APEX2* (Alvarez-Quilon *et al*, 2020; Fleury *et al*., 2022; Mengwasser *et al*., 2019) (Fig. S1F). Notably, we retrieved all members of the 9-1-1 complex (RAD9A-HUS1-RAD1) and its interacting partner RHINO (encoded by *RHNO1*) as essential in TKO cells (Fig. 1d-e). To confirm the synthetic lethality observed in our screen, we individually targeted subunits of the complex using two independent sgRNAs and performed cellular growth assays (Fig. 1f, S1G-I). While depletion of 9-1-1 and RHINO had little impact on control cells, their loss significantly compromised the survival of TKO cells (Fig. 1f), thereby validating their roles in promoting the survival of cells defective in HR and NHEJ.

### RAD9-HUS1-RAD1 (9-1-1) and RHINO are critical MMEJ factors

To directly test whether the 9-1-1 complex and RHINO are involved in MMEJ-mediated repair, we conducted experiments to assess the repair of dysfunctional telomeres in cells lacking the shelterin complex. The six-subunit shelterin complex protects telomeres from the DNA damage response. When shelterin is absent, telomeres become deprotected and activate a DNA damage response at chromosome ends (Sfeir & de Lange, 2012). In cells lacking the NHEJ factors Ku70/80, MMEJ is the primary repair pathway at deprotected telomeres, leading to chromosome end-fusions in a POLQ, LIG3, and PARP1-dependent manner (Sfeir & de Lange, 2012) (Fig. 2a). To determine the role of the 9-1-1 complex and RHINO in MMEJ, we individually targeted these factors in TRF1/2^Δ^Ku80^-/-^ mouse embryonic fibroblasts (MEFs) (Fig. S2A-B) and noted a significant reduction in the frequency of telomere fusions (Fig. 2b-c). However, depletion of the 9-1-1 subunits or RHINO in Ku80^+/+^ MEFs, where NHEJ drives telomere fusions, did not affect telomere fusions, suggesting that the activity of 9-1-1 and RHINO is specific to repair by MMEJ (Fig. 2d).

**Figure. 2.**
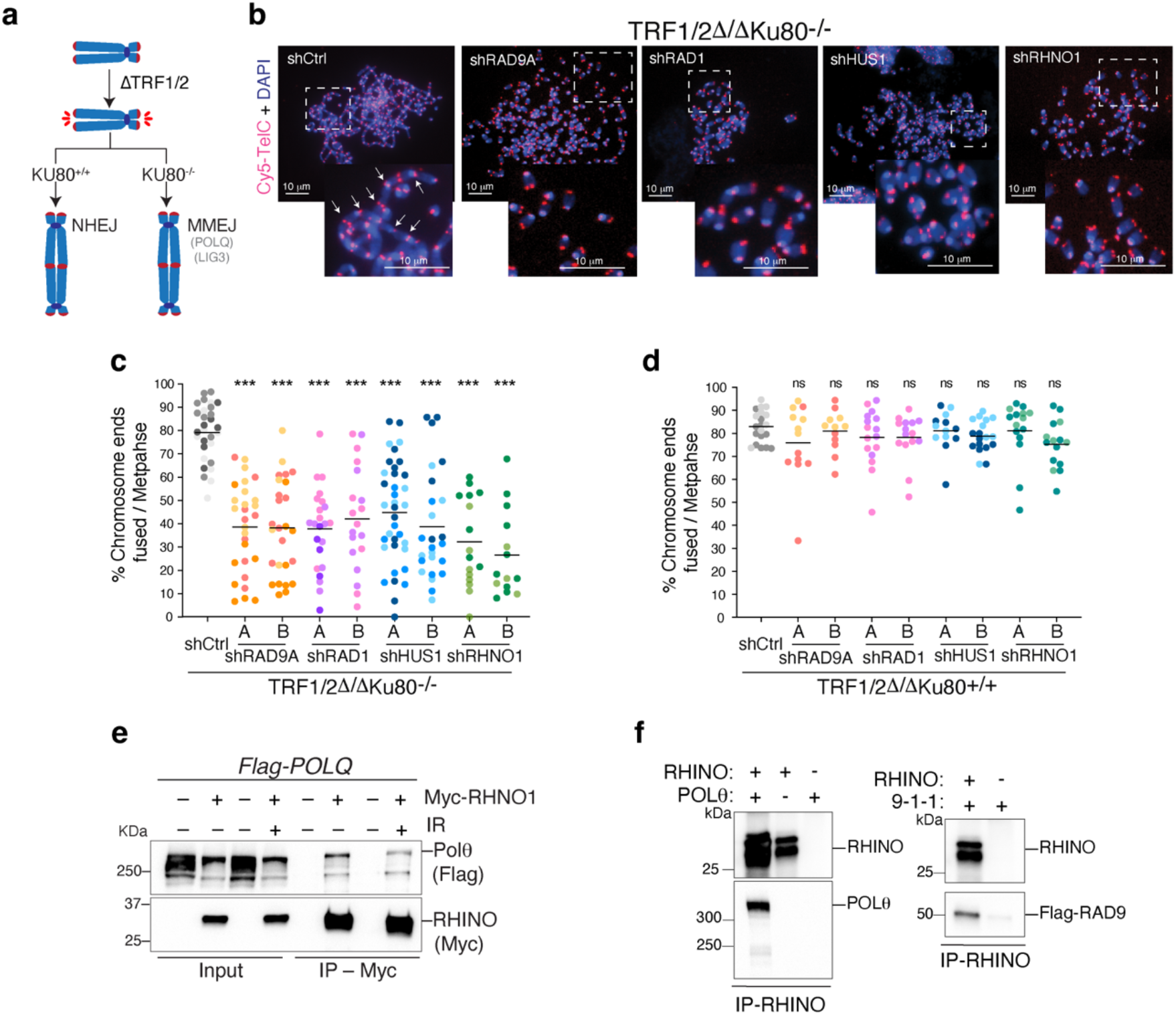
A non-canonical function for 9-1-1/RHINO in MMEJ. (**A**) Schematic representation of the shelterin-free assay (Sfeir & de Lange, 2012) to monitor MMEJ frequency at deprotected telomeres. Deletion of TRF1 and TRF2 cause the loss of the remaining shelterin subunits from telomeres and results in chromosome end-fusions by NHEJ (Ku80^+/+^) and MMEJ (Ku80^-/-^). (**B**) Representative images of a metaphase spread from TRF1/2^Δ/Δ^Ku80^−/−^p53^−/−^ cells depleted for individual subunits of the 9-1-1 complex or RHINO. Telomeres are marked by FISH using a Cy5-[CCCTAA]3 PNA probe (red), and chromosomes are counterstained with DAPI (blue). White arrows indicate examples of telomeric fusions in the control sample. (**C**) Quantification of telomere fusions mediated by MMEJ related to panel b. Data are the mean of at least two independent experiments. The different colors used for the dots represent independent experiments. (**D**) Quantification of telomere fusions by NHEJ in TRF1/2^Δ/Δ^Ku80^+/+^. Data are the mean of at least two independent experiments. The different colors used for the dots represent independent experiments. (**E**) Co-immunoprecipitation (Co-IP) experiments depicting Polθ and RHINO interaction in lysates from whole-cell extracts from HEK293T cells co-transfected with plasmids expressing FLAG-Polθ and MYC-RHINO. Co-IPs were performed on cells treated with ionizing radiation (20 Gy) and control cells. (**F**) *In vitro* Co-IP experiments showing Polθ/RHINO and RAD9/RHINO interaction with purified proteins. Polθ was purified from HEK293T cells, RHINO from *E. coli*, and RAD9 from *S. cerevisiae*.

We next evaluated the impact of 9-1-1 and RHINO depletion on DNA damage signaling by examining the co-localization of the DNA damage marker 53BP1 with telomeric DNA. Even though there was a significant decrease in MMEJ activity, the absence of 9-1-1 and RHINO in shelterin-free cells did not hinder the accumulation of 53BP1 at deprotected telomeres (Fig. S2C-D). This indicates that the canonical role of the 9-1-1 complex in activating ATR signaling alone may not be solely responsible for its role in MMEJ, and other mechanisms may be involved.

In an orthogonal approach, we measured MMEJ activities at an I-SceI-induced break using the traffic light reporter (TLR) system (Certo *et al*, 2011) (Fig. S2E). To limit repair to MMEJ and HR, we stably expressed the TLR reporter in *LIG4*^*-/-*^ cells where NHEJ is blocked. We first treated these cells with sgRNA targeting 9-1-1 and RHINO, then introduced I-SceI to trigger break formation. The percentages of mCherry and GFP-positive cells were then quantified to reflect MMEJ and HR activities, respectively. Consistent with a critical role for the 9-1-1 complex and RHINO during MMEJ repair, we observed a significant reduction in the percentage of mCherry by TLR in cells depleted for subunits of the 9-1-1 complex and RHINO (Fig. S2F-I). The results from the TLR assay further confirm the telomere fusion assay (Fig 2b-c) and are consistent with Repair-seq data, which revealed a correlation between subunits of the 9-1-1 complex and *POLQ* (Hussmann *et al*, 2021).

### A non-canonical function for 9-1-1 and RHINO in MMEJ

The heterotrimeric 9-1-1 complex is loaded onto 5’ ends of resected DNA and single-stranded DNA gaps in response to replication stress (Melo *et al*, 2001). Its interaction with RHINO was reported to amplify cell cycle checkpoint signaling through the ATR kinase (Cotta-Ramusino *et al*, 2011; Lindsey-Boltz *et al*, 2015; Mordes *et al*, 2008). However, ATR is also activated independently of the 9-1-1/TOPBP1 axis through ETAA1/ATRIP, which binds directly to RPA and is recruited to DNA damage sites (Bass *et al*, 2016). Interestingly, the synthetic lethal screen did not identify hits for *ATRIP* or *ETAA1* (Fig. 1e). Deleting these genes using CRISPR/Cas9 did not lead to growth defects in TKO cells (Fig. 1f, S1G-I). Moreover, the depletion of ETAA1 and ATRIP in TRF1/2^Δ*/*Δ^Ku80^-/-^ MEFs only mildly affected telomere fusions (Fig. S3A-E). This contrasts the significant impact on telomere fusion observed with 9-1-1, RHINO, and POLQ depletion (Fig. S3A-C). In conclusion, our results show that inhibition of ATR signaling by depletion of ATRIP and ETAA1 was insufficient to block MMEJ. These data further suggest that the role of 9-1-1 and RHINO in ATR signaling is insufficient to promote MMEJ.

The 9-1-1 complex forms a ring shape that bears a striking structural resemblance to proliferating cell nuclear antigen (PCNA), another DNA-encircling complex that recruits and physically interacts with various DNA polymerases during replication (Eichinger & Jentsch, 2011). We suspected that 9-1-1 and RHINO could foster MMEJ by interacting with Polθ and recruiting it to break sites. To test this, we performed Co-IP experiments in HEK293T cells by co-expressing 9-1-1 complex proteins and RHINO with Flag-tagged POLQ. While we could not detect an interaction between Polθ and any of the subunits of the 9-1-1 complex (Fig. S3F), we provided evidence for Polθ interaction with RHINO, independent of DNA damage (Fig. 2e). In addition, we probed for a direct interaction between RHINO and Polθ by performing Co-IP using purified proteins (Fig. S3G) and demonstrated that RHINO directly binds full-length Polθ (Fig. 2f). Collectively, our data implicate the RHINO-Polθ interaction in promoting MMEJ.

### RHINO expression is confined to mitosis

RHINO was initially identified in an RNAi screen designed to identify factors that promoted cell cycle arrest following irradiation (Cotta-Ramusino *et al*., 2011). To gain better insight into the function of RHINO beyond checkpoint signaling, we sought to determine its genetic interactors and highlight pathways essential in RHINO-deficient cells. To that end, we carried out CRISPR/Cas9 synthetic lethal screens in three clonally derived RHNO1^-/-^ cells and the parental RHNO1^+/+^ cell line (Fig. S4A-E). As expected, with MMEJ deficiency, HR factors, including FANCD2 and BRCA2, were essential for RHNO1^-/-^ cell growth (Fig. 3a, S4F). Notably, mitosis-related genes were among the top hits, including cyclin-dependent kinase CDK1, the spindle checkpoint proteins *BUB3* and *BUB1B*, and the kinetochore factors *ZWILCH* and NDC80 (Fig. 3a). Furthermore, pathway analysis of genes essential in RHNO1^-/-^ cells revealed the enrichment in several pathways related to mitosis (Fig. 3b). Independently, we analyzed the genetic dependencies in the DepMap database (Institute, 2021) and found that *RHNO1* correlated with *CIP2A, MDC1*, and *TOPBP1*, which forms a complex that tethers mitotic DSBs together (Adam *et al*., 2021; De Marco Zompit *et al*, 2022; Leimbacher *et al*, 2019). DepMap analysis also uncovered a correlation between the essentiality scores of *POLQ, CIP2A*, and *RHNO1* (Fig. 3c).

**Figure. 3.**
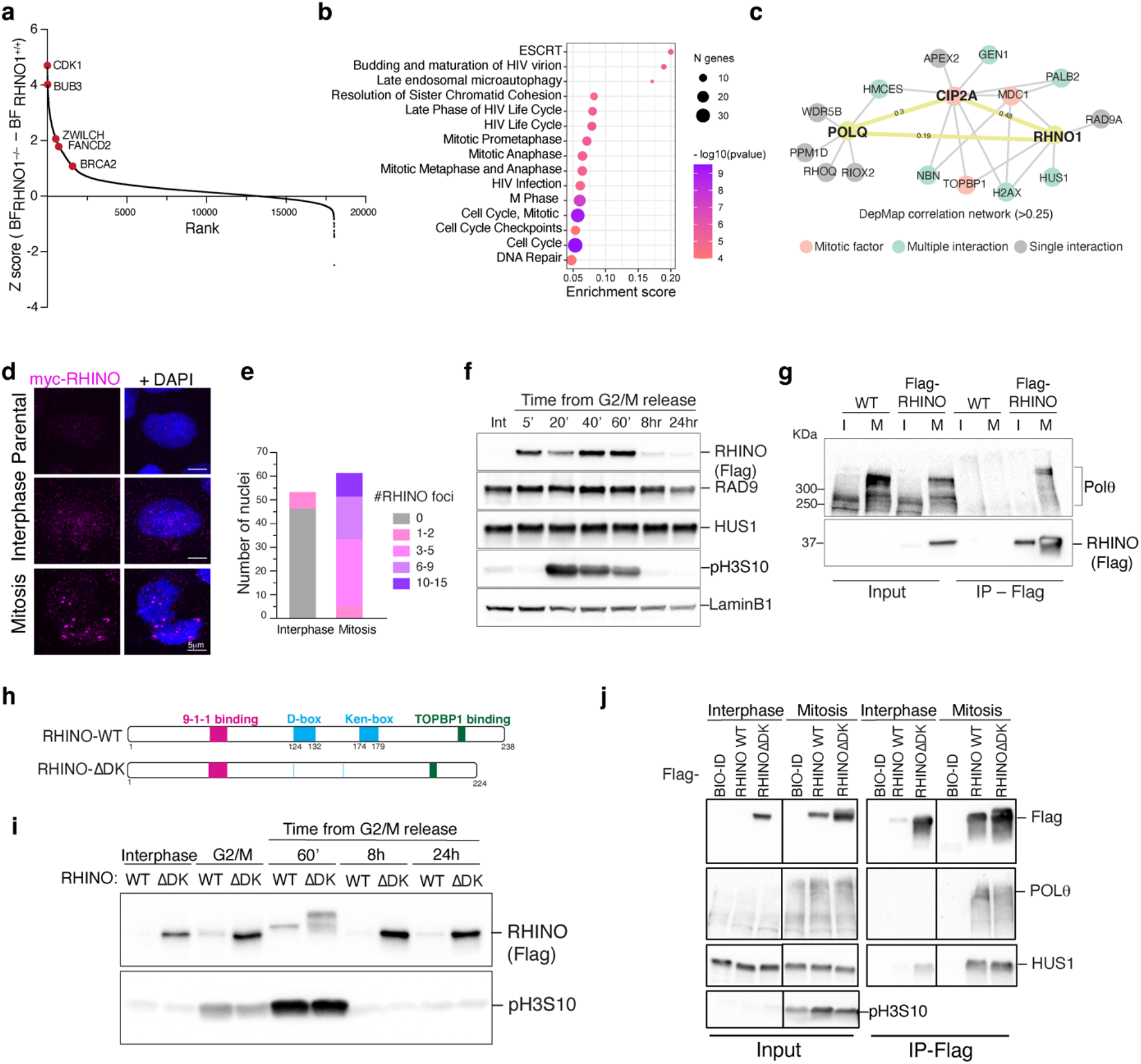
RHINO expression is confined to mitosis. (**A**) Results from a CRISPR/Cas9 dropout screen in *RHNO1*^*-/-*^ and isogenic *RHNO1*^*+/+*^ DLD1 cells. Ranked z-scores of the difference in Bayes factor (BF) scores for essential genes in *RHNO1*^*-/-*^ *vs. RHNO1*^*+/+*^. BF scores were determined using the BAGEL2 analysis pipeline. Highlighted in red are HR factors, including *BRCA2* and *FANCD2*, and mitotic factors *CDK1, BUB3*, and *ZWILCH*. The dataset is available in Supplementary Table 1. (**B**) Reactome pathway overrepresentation analysis of synthetic lethal genes with *RHNO1*^*-/-*^. Fold enrichment of each pathway is plotted on the X-axis. The number of genes associated with each pathway is indicated by the size of the circle, while the color shade of the circles indicates the p-value. (**C**) Network analysis for POLQ and *RHNO1* based on Pearson’s correlation of dependency scores derived from DepMap. (**D**) Representative immunofluorescence images of DLD1 cells stably overexpressing FLAG-MYC-RHINO during interphase and mitosis stained with anti-Myc antibody. (**E**) Quantifying RHINO foci in mitosis and interphase as in panel D. (**F)** Western blot analysis of 9-1-1/RHINO expression during the cell cycle. Cells stably overexpressing FLAG-MYC-RHINO were arrested in G1/S using a double thymidine block and then released in CDK1 inhibitor for 16h to induce a G2/M arrest. Cells were collected at the indicated time points and subjected to Western blot to detect FLAG-MYC-RHINO and endogenous components of the 9-1-1 complex. The phosphoantibody against serine 10 in H3 (pS10H3) was used as a mitotic marker. Lamin B1 was used as a loading control. Int = interphase. (**G**) Cells overexpressing MYC-FLAG-RHINO^ΔDK^ and WT were synchronized in mitosis and subjected to immunoprecipitation followed by Western blot for endogenous Polθ. I = interphase M = mitosis. (**H**) Schematic of RHINO^ΔDK^ mutant. (**I**) Cells overexpressing MYC-FLAG-RHINO and MYC-FLAG-RHINO^ΔDK^ were synchronized using a double thymidine block followed by CDK1 inhibitor and released with fresh media into mitosis. Samples were taken at the indicated time points after the G2/M release. “G2/M” indicates cells harvested without washing out the CDK1 inhibitor. 30ug of cell lysate was subjected to Western blot to detect FLAG-RHINO, FLAG-RHINO^ΔDK,^ and pS10H3 as a mitosis marker. (**J**) Cells overexpressing FLAG-RHINO, FLAG-RHINO^ΔDK^, and FLAG-BioID were synchronized using a double thymidine block followed by CDK1 inhibition and released into nocodazole to enrich mitotic cells. “Interphase” indicates cells that did not undergo any synchronization. FLAG-BioID is used as a negative control. Cell lysates were subjected to anti-FLAG immunoprecipitation. 30ug of protein was loaded as input, and 25% of the immuno-antigen eluant was loaded as the IP fraction on SDS-PAGE gels and then subjected to Western blot using the indicated antibodies.

The synthetic lethal screen and DepMap analysis results uncover a previously unknown role for RHINO in mitosis. This is supported by the observation that RHINO displayed large foci in cells arrested in M phase and visualized using immunofluorescence (Fig. 3d-e). Interestingly, Western blot analysis on samples taken at different cell cycle stages demonstrated that RHINO expression is restricted to mitosis (Fig. 3f, S4G-H), where it gets phosphorylated irrespective of DNA damage (Fig. S4I-J). In contrast, subunits of the 9-1-1 complex were ubiquitously expressed throughout the cell cycle (Fig. 3f, S4G-H). As predicted by the strict expression of RHINO, the interaction between stably expressed Flag-RHINO and endogenous Polθ was exclusive to mitosis (Fig. 3g).

RHINO protein contains two recognition sequences for the Anaphase-promoting complex (APC/C), namely the Ken-box and D-box domains (Fig. 3h). To investigate whether APC/C played a role in the degradation of RHINO during mitotic exit, we stably expressed RHINO^ΔDK^ that carries deletions in both degrons. We assessed RHINO expression throughout the cell cycle. Blocking APC/C-mediated ubiquitination and subsequent degradation resulted in RHINO stabilization beyond mitosis (Fig. 3i). Despite being stable in interphase, RHINOΔDK interacted with Polθ exclusively in mitosis (Fig. 3j), suggesting that post-translational modifications, such as phosphorylation by mitotic kinases, may be necessary for the RHINO-Polθ interaction. In conclusion, these findings provide new insights into the temporal regulation of MMEJ and demonstrate a previously unanticipated regulation of RHINO during the cell cycle. Specifically, RHINO accumulates exclusively in mitosis, interacts with Polθ, and is rapidly degraded upon mitotic exit by APC/C. The stabilization of RHINO upon mitotic entry remains unclear and warrants further investigation.

### MMEJ is the dominant DSB repair pathway in mitosis

Based on the critical function of RHINO in MMEJ and its exclusive expression in mitosis, we hypothesized that MMEJ might repair breaks specifically in M phase, where NHEJ and HR are repressed (Blackford & Stucki, 2020). Our hypothesis was supported by circumstantial evidence from the literature implicating Polθ activity in chromosome rearrangements in mitosis (Anne Margriet Heijink *et al*, 2021; Deng *et al*, 2019; Llorens-Agost *et al*, 2021; Wang *et al*, 2018). To provide direct evidence that links MMEJ activity to mitosis, we synchronized *POLQ*^*-/-*^ and *POLQ*^*+/+*^ cells at the G2/M boundary using a CDK1 inhibitor (RO3306) and then released cells in the presence of nocodazole, preventing mitotic exit. Cells were irradiated 30 minutes after release from CDK1 inhibition, and γH2AX levels were measured at one- and five-hours post-irradiation (Fig. 4a). Wild-type cells accumulated maximal H2AX foci intensity one-hour post-irradiation that was significantly H2AX foci intensity one-hour post-irradiation that was significantly decreased after five hours (Fig. 4b-c, S5A). In contrast, *POLQ*^*-/-*^ cells failed to resolve γH2AX foci (Fig. 4b-c). Cells carrying inactivating mutations in the polymerase domain of Polθ (*POLQ*^*ΔPol/ΔPol*^) behaved similarly to *POLQ*-deficient cells (Fig. 4B-C). We confirmed the primary role of MMEJ in repairing DSBs in mitosis by treating cells with RP6685, a Polθ polymerase inhibitor (Monica Bubenik*, 2022), that significantly impaired the resolution of DNA damage and resulted in the retention of γH2AX foci at 5 hours post-irradiation (Fig. S5B). In addition, we assessed the impact of mitotic MMEJ on cellular growth. We showed that *POLQ*^*-/-*^ cells were more sensitive to DNA damage by ionizing radiation in mitosis than control cells (Fig. S5C-D). To test if mitotic MMEJ can repair DNA lesions that arise in S and G2 phases and persist into M phase, we employed a strategy to enrich for unrepaired breaks in S phase (Fig. 4d). First, we treated cells with siRNA against *BRCA2* to block repair by HR (Fig. S5E). We then arrested siRNA-treated cells at the G1/S boundary using thymidine.

**Figure. 4.**
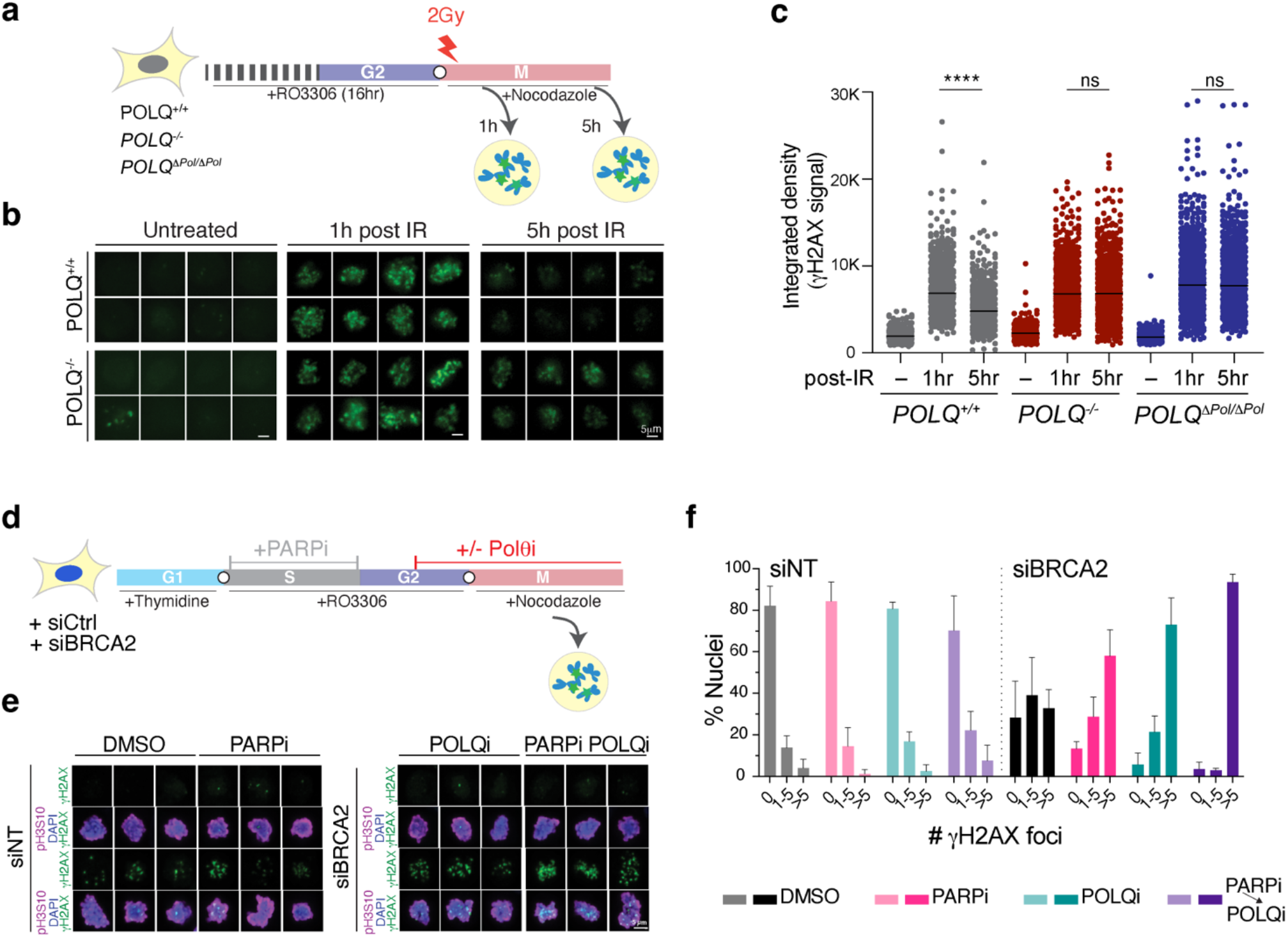
MMEJ is the predominant DSB repair pathway during mitosis. (**A**) Schematic of the experimental pipeline for panels B and C. Cells are synchronized at the G2/M boundary with the CDK1 inhibitor RO3306 for 16h and then released into mitosis in nocodazole. Cells are irradiated with 2 Gy 30 minutes after release into mitosis and fixed for immunofluorescence (IF) with γH2AX antibody at one and five hours after irradiation. (**B**) Representative images of γH2AX stain (Green) in cells treated as described in panel A. (**C)** Quantification of γH2AX intensity in mitotic cells in parental, *POLQ* knockouts or cells with DNA polymerase catalytic-dead *POLQ* mutant (*D2540A, E2541A*). Statistical analysis was performed using a non-parametric unpaired t-test (****p<0.0001). At least 450 nuclei per sample were imaged for the experiment. (**D**) Schematic of the experimental pipeline for panels E-F. Cells were synchronized at the G1/S interphase using thymidine block and released into S-phase in the presence of both PARP inhibitor (Olaparib) and CDK1 inhibitor (RO3306). PARP inhibitor was withdrawn upon exit from S phase and replaced with a Polθ inhibitor at the end of G2 and into M phases. Cells were treated with nocodazole, fixed, and stained with γH2AX antibody one hour after release into mitosis. (**E**) Representative immunofluorescence images of mitotic cells treated as described in panel D and stained with anti-γH2AX. pH3S10 is used to stain mitotic chromosomes (Magenta). DNA is stained in blue with DAPI. (**F**) Quantification of γH2AX foci in mitotic cells with the indicated treatment. At least 50 nuclei were counted for the experiment. Bars represent the mean of 3 independent experiments.

We released them into S phase in the presence of both CDK1 inhibitors to prevent entry into mitosis and PARP inhibitor (Olaparib – 1μM) to induce S phase damage (Murai *et al*, 2012). As cells exited S phase, we withdrew Olaparib and added Polθ inhibitor (RP6685 – 10μM) towards the end of G2. We maintained the Polθ inhibitor after swapping the CDK1 inhibitor with nocodazole to prevent exit from mitosis (Fig. 4d and S5E-F). We observed a baseline increase in the number of γH2AX foci in mitotic cells treated with siBRCA2 compared to those treated with control siRNA. As expected, the levels of γH2AX were even more significant in cells treated with either Olaparib in S phase or RP6685 in G2/M phase (Fig. 4e-f, S5G). Strikingly, cells treated with both PARP inhibitor in S phase and Polθ inhibitor in G2/M displayed a synergistic increase in the levels of γH2AX foci (Fig. 4e-f, S5G). In an independent experiment, we treated cells with Polθ inhibitor only in mitosis and observed a similar synergy with PARP inhibitor treatment in S phase (Fig. S5H). These results provide direct evidence that MMEJ is the predominant DSB repair pathway in mitosis, fixing damage incurred in M phase and unrepaired breaks originating in S phase.

### RHINO recruits Polθ to DSB sites in mitosis

Having established a role for MMEJ in repairing breaks in mitosis, we next aimed to investigate if RHINO recruits Polθ to mitotic damage sites. To achieve this, we utilized a two-step CRISPR/Cas9 targeting strategy to introduce a Halo tag at the N-terminus of *POLQ*, enabling us to visualize endogenous Polθ in live cells (Fig. S6A-C). We labeled *POLQ*^*Halo/Halo*^ with a Halo-tag ligand (JFX650) and traced Polθ single-particles in live-cell imaging in interphase and mitosis (Fig. S6D and Supplemental movies 1-2). To test if RHINO acts upstream of Polθ, we arrested *POLQ*^*Halo/Halo*^ cells at the G2/M boundary by CDK1 inhibition. We released them into mitosis in the presence of the DNA damage agent zeocin. This resulted in the appearance of large and static Polθ foci that were reduced upon RHNO1 depletion using two different siRNAs (Fig. 5a-b and S6E).

**Figure. 5.**
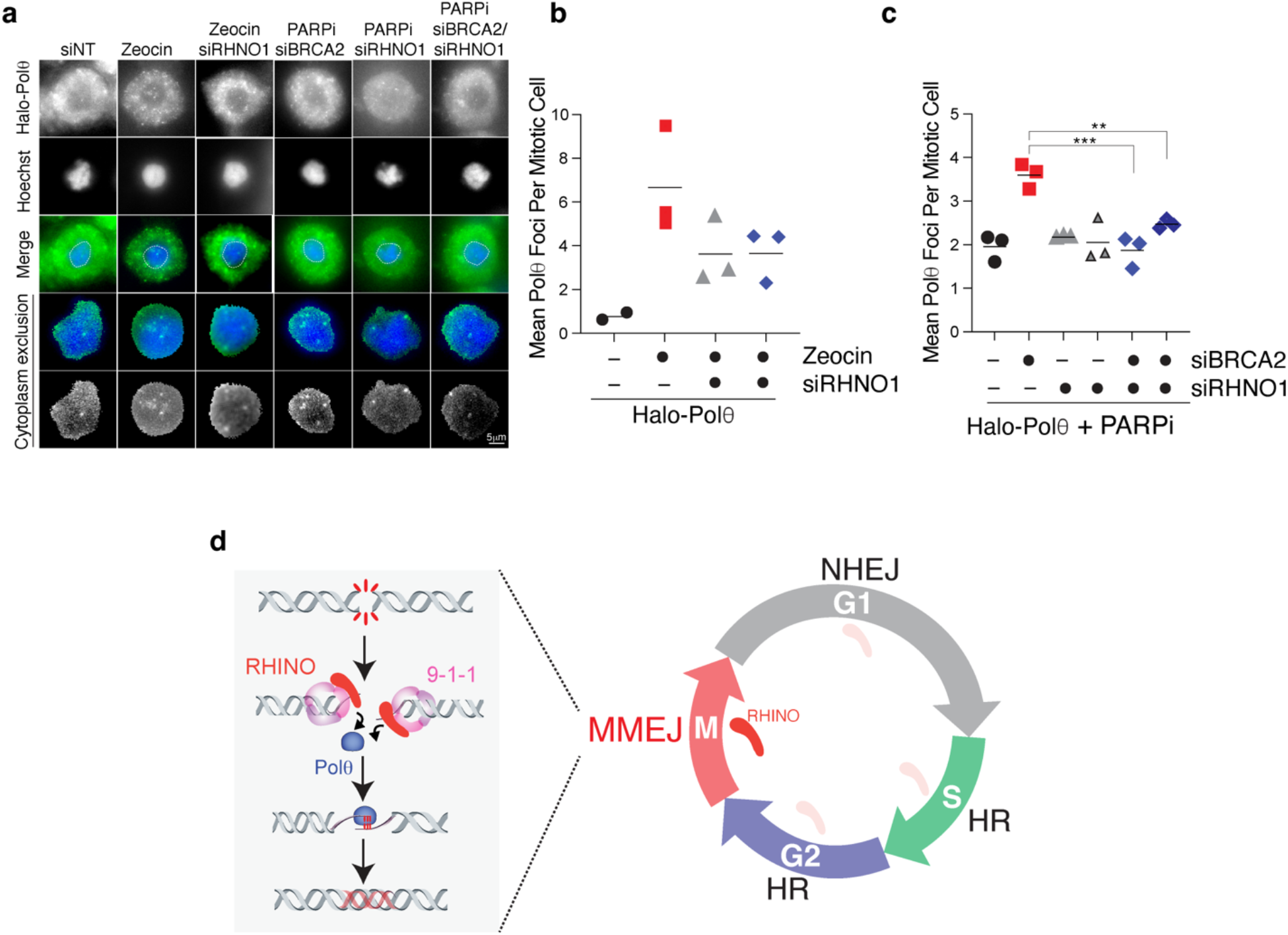
RHINO recruits Polθ to damaged sites in mitosis. (**A**) Representative images of Halo-Polθ foci in mitosis. Cells were transfected with the indicated siRNA and synchronized at the G1/S interphase using a thymidine block and released into S-phase in the presence of a CDK1 inhibitor (RO3306). Cells were arrested in mitosis with nocodazole, similarly to Fig. 4D. For Zeocin treatment, cells were arrested in mitosis with nocodazole and treated with Zeocin (20 μg/mL; 1 hour). For experiments assessing Polθ foci formation in mitotic cells with persistent damage from S phase, cells expressing siRNA against *BRCA2* and *RHNO1* were treated with Olaparib during S phase according to the schematic in Fig. 4D. Scale bar = 5 μm. (**B**) Quantification of mitotic Halo-Polθ foci in live cells treated with Zeocin and depleted of *RHNO1*. Bars represent the mean of 3 independent experiments. At least 40 nuclei were counted for the experiments. (**C**) Quantification of mitotic Halo-Polθ foci in live imaging experiments in nocodazole arrested cells treated with PARP inhibitor during S-phase. The cells were treated as described in panel A. Bars represents the mean of 3 independent experiments. Statistical analysis was performed using one-way ANOVA (***p<0.001, **p<0.01). At least 40 nuclei were counted for these experiments. (**D**) Model for RHINO restricting MMEJ activity to mitosis. Uncoupling of DNA repair pathways as a function of the cell cycle. NHEJ predominates in G1. HR is the preferred pathway for S and G2. MMEJ is the only DSB repair pathway in mitosis. The tight expression of RHINO, which recruits Polθ to break sites in mitosis, restricts MMEJ to M phase.

Last, we investigated the static Polθ foci in the context of unresolved S phase damage that persisted into mitosis. We treated *POLQ*^Halo/Halo^ cells with siRNA against BRCA2 (Fig. S6E) and PARP inhibitor (Olaparib– 1μM) in S phase and monitored Polθ foci formation in live cells. This treatment did not alter Polθ dynamics or lead to any detectable co-localization of Polθ with RPA foci in S phase (Fig S6F). However, when BRCA2 was depleted in cells and treated with PARP inhibitors in S phase, static Polθ foci accumulated in mitosis and frequently co-localized with RPA (Fig. 5a and 5c and S6G-H). Importantly, we found that the accumulation of Polθ in mitosis was dependent on RHINO (Fig. 5a and 5c). Taken together, these data further implicate RHINO in recruiting Polθ to break sites in mitosis to promote MMEJ at unrepaired lesions.

## Discussion

Condensed mitotic chromosomes are highly vulnerable to DNA damage (Zirkle & Bloom, 1953), and it is well established that both HR and NHEJ activities are blocked in mitosis (Blackford & Stucki, 2020; Rieder & Cole, 1998). While the initial detection of breaks and the accumulation of γH2AX in mitotic cells remain intact (Giunta *et al*, 2010), recruitment of RNF8 and 53BP1 is impaired, leading to inhibition of downstream signaling and suppression of NHEJ (Giunta *et al*., 2010; Nelson *et al*, 2009). Similarly, HR factors, including *BRCA1/2* and *RAD51*, do not engage with mitotic DSBs (Ayoub *et al*, 2009; Giunta *et al*., 2010). Unrepaired mitotic breaks are highly deleterious and can lead to lagging chromosome fragments, chromosome missegregation, and accumulation of micronuclei (Crasta *et al*, 2012). Therefore, cells must have evolved mechanisms to safeguard their genetic material during this critical time in the cell cycle. Recent studies proposed that cells circumvent mitotic problems due to DNA breaks via the MDC1-TOPBP1-CIP2A complex, which tethers broken ends of mitotic DNA until cells exit mitosis and enter G1, where NHEJ becomes active (Adam *et al*., 2021; De Marco Zompit *et al*., 2022). Here, we provide direct evidence for a robust mitotic DNA repair activity by MMEJ and uncover a critical role for RHINO in directing this error-prone repair activity to M phase (Fig. 5d). Our findings are consistent with previous findings implicating Polθ activity in mitosis. This includes a study using *Xenopus* egg extract, showing that entry into mitosis before completion of DNA replication leads to complex rearrangements driven by Polθ (Deng *et al*., 2019). In addition, Polθ was recently linked to the formation of Sister Chromatid Exchanges (SCE) as under-replicated DNA is transferred into mitosis (Anne Margriet Heijink *et al*., 2021). Last, our data aligns with recent work showing that BRCA2 and Rad52 prevent Polθ activity in S and G2 (Llorens-Agost *et al*., 2021). Further studies are needed to elucidate the interplay between DSB repair by Polθ-mediated MMEJ and DSB tethering by MDC1-TOPBP1-CIP2A.

Based on our results, we provide a conceptually novel framework for DSB repair as a cell cycle function, with NHEJ predominating in G1, HR being the preferred pathway for S and G2, and MMEJ being the pathway of choice for mitosis (Fig. 5d). The uncoupling of DNA repair pathways during different stages of the cell cycle has significant implications for maintaining genome stability. The employment of MMEJ in mitosis may have evolved as a failsafe mechanism to ensure that cells do not commit to cellular division with unrepaired lesions. On the other hand, suppression of MMEJ in G1 and S/G2 could protect our genome from the intrinsic mutagenicity of this pathway.

Our study identifies a novel function for RHINO as a mediator of Polθ recruitment to break sites in mitosis to promote MMEJ. We cannot rule out that 9-1-1 and RHINO foster MMEJ by amplifying ATR signaling in mitosis. However, Co-IP analyses using lysates and purified proteins reveal a direct RHINO-Polθ interaction (Fig. 2e-f). Furthermore, we show that inhibition of ETAA and ATRIP does not impact MMEJ (Fig. 1f, S3A-E). Last, 9-1-1 and RHINO depletion prevent chromosome fusions by MMEJ, despite the persistence of DNA damage signaling at deprotected telomeres. Based on these results, we favor a model where RHINO stimulates MMEJ by recruiting Polθ to break sites. The role of RHINO in MMEJ is consistent with its original identification in a siRNA screen for DNA damage factors that sensitize cells to irradiation (Cotta-Ramusino *et al*., 2011). Our data demonstrate that RHINO, but not 9-1-1 complex members, is strictly expressed in mitosis. Its expression is tightly regulated by targeted depletion by the APC/C complex (Fig. 3h-i). RHINO is an unstructured protein interacting with RAD1 and TOPBP1 using separate domains (Cotta-Ramusino *et al*., 2011; Lindsey-Boltz *et al*., 2015). One possible scenario is that unrepaired S phase damage is marked by 9-1-1 until the onset of mitosis when RHINO accumulates. Tethering RHINO to the 9-1-1 complex in M phase would subsequently lead to Polθ recruitment to break sites and promote MMEJ. It is also plausible that a RHINO-TOPBP1 interaction in mitosis could stabilize RHINO at break sites. Finally, our data do not rule out the possibility of additional roles for Polθ in filling ssDNA gaps accumulating during S phase and persisting through mitosis (Belan *et al*, 2022; Mann *et al*, 2022; Roerink *et al*, 2014; Schrempf *et al*, 2022). In conclusion, our study provides evidence suggesting robust MMEJ activity in mitosis that may account for the synthetic lethal interaction between Polθ and BRCA2. Furthermore, uncoupling DNA repair activities during different cell cycle stages provides a rationale for the reported synergy of Polθ inhibitors with PARP inhibitors and potentially other antineoplastic therapies that induce DNA damage during S-phase (Zatreanu *et al*., 2021; Zhou *et al*., 2021).

## Acknowledgments

We thank Dan Durocher for providing critical reagents and Robert Weiss for kindly sharing Rad9, Rad1, and Hus1 expression plasmids. We thank Stephen Morris, Karl Zahn, and Marie-Claude Mathieu for providing full length Polθ protein and Dirk Remus for providing 9-1-1 proteins and helping with *in vitro* immunoprecipitation experiments. We thank Eros Lazzerini-Denchi and members of the Sfeir lab for commenting on the manuscript. We thank Sarah Deng for generating Lig4^-/-^ 293T cells. This work was supported by a grant from the NIH /NCI (R01 CA229161) to A.S. and from the Pershing Square-Sohn and the V-foundation (A.S.). A training grant from NYSTEM supported A.B. J.H. is supported by an NIH fellowship (F32GM139292). A DP2 from the NIH (DP2GM142307) supports work in the J.S. lab.

## Authors Contribution

A.S. and A.B. conceived the experimental design. A.B. performed or oversaw all experiments. S.P. and A.B. performed experiments reported in Fig. 4. M.K. helped analyze telomere fusions reported in Fig. 2. O.S. performed experiments related to RHINO-PolQ interaction and RHINO expression in mitosis (Fig. 2 and Fig. 3). J.H., in the lab of J.S., performed live-cell imaging of Polθ in mitosis (Fig 5). A.S. and A.B. wrote the manuscript. All authors discussed the results and commented on the manuscript.

## Methods

### Cell culture

UO2S and HEK293T cells were purchased from ATCC and grown in Dulbecco’s Modified Eagle Medium (DMEM; Gibco/Thermo Fisher) supplemented with 10% fetal bovine serum (FBS, Gibco/Thermo Fisher). *TRF1/2*^*Δ/Δ*^*Ku80*^*−/−*^*p53*^*−/−*^ and *TRF1/2*^*Δ/Δ*^ *Ku80*^*+/+*^*p53*^*−/−*^ MEFs were previously established (Sfeir & de Lange, 2012). Parental and *BRCA2*^*-/-*^ DLD-1 cells were purchased from ATCC (CCL-221) and Horizon, respectively (HD105-007) and maintained in RPMI medium (Gibco/Thermo Fisher) supplemented with 10% FBS. Cells expressing Halo-Polθ were grown in RPMI medium supplemented with 10% FBS. All cells were supplemented with 0.1 mM MEM non-essential amino acids (Gibco/Thermo Fisher), 2 mM L-glutamine, 100 U/ml penicillin and 100 μg/ml streptomycin (Pen/Strep; Gibco/Thermo Fisher) and grown at 37 °C and 5% CO_2._ To synchronize in G1, cells were treated for 18-24h with 2 mM of thymidine (MilliporeSigma, T1895-5G) followed by 16h of incubation with 9 μM of RO-3306 (SelleckChem, catalog no. S7747) to synchronize at the G2/M phase. To allow cells to enter and accumulate in mitosis, cells were released from G2/M blockade in 100 ng/ml of Nocodazole (MilliporeSigma, M1404). In experiments with PARP inhibitors, we used Olaparib at 1 μM or 10 μM as indicated in figure legend for DLD1 and U2OS (Selleckchem, S1060). For Zeocin-treated samples, cells were treated for one hour with 20 μg/mL Zeocin (ThermoFisher R25005). Cells were irradiated with X-RAD225 - Precision X-Ray – (Accela).

### Plasmids

The gRNAs used to knock-out *LIG4* were cloned in the pSpCas9(BB)-2A-GFP (pX458) (Addgene #48138). The gRNA used for IncucyteS3 experiments was cloned into a pLenti-gRNA-GFP-2A-PURO gift from Jordan Young from Repare Therapeutics. The lentiviral Cas9-BLAST was purchased from Addgene (#52962). The shRNA that knocked down *9-1-1/RHNO1, ETAA1* and *ATRIP* in MEFs was cloned into the pLKO.1 vector from the RNAi consortium (Broad Institute). The 9-1-1 plasmids were a gift of Robert Weiss (pCMV-RAD1-HA, pCMV14-HUS1-3XFLAG, pCMV-RAD9A-MYC). A lentiviral RHINO-myc-Flag plasmid (pCMV6-RHNO1-Myc-FLAG) was purchased from Origene (RC203020). Full-length human *POLQ* cloned into pLPC-Flag vector was previously described (Mateos-Gomez *et al*., 2015). To tag endogenous *POLQ* at the N-terminus with a 3xFLAG-LoxP-SV40-Puro-Lox-HaloTag (Addgene #86843), ∼ 5 × 10^5^ U2OS cells were transfected using FuGene6 (Promega) with 1μg of gRNA/Cas9 (px330) and HDR plasmid in 6 well plates. Edited cells were selected with 1 μg/mL of puromycin. Single-cell clones were sorted, and N-terminally edited cells were transfected with a plasmid encoding Cre to recombine the PuroR cassette generating a 3xFLAG-HaloTagged-*POLQ* protein. Cells were labeled with 250 nM JF646-HaloTag ligand (JF646) sorted based on the JF646 signal (Grimm *et al*, 2015). Homozygous clones were identified by genomic PCR using primers orientated outside each homology arm. To visualize RPA in live imaging experiments, ∼ 6 × 10^5^ U2OS cells were transfected with 1.8μg of a mNEON-RPA32 plasmid.

### Antibodies

Primary antibodies used for Western blot included FLAG (Clone M2, Sigma; 1: 10000 dilution), MYC (9B11; Cell Signaling; 1: 1000 dilution), LIG4 (HPA001334; Sigma; 1: 500 dilution), p53 (sc126; Santa Cruz; 1: 1000 dilution), RAD1 (NB100-346; Novus Biologicals; 1: 1000 dilution), RAD9 (611324, BD Transduction Laboratories, 1: 500 dilution), HUS1 (D4J9H, Cell Signaling Technology; 1: 1000 dilution), Polθ (gift from Repare therapeutics; 1: 1000 dilution), phospho-H2AX (S139, 9718 S; Cell Signaling Technology; 1: 1000 dilution), phospho-H3 (S10, 9718 S; Cell Signaling Technology; 1: 10000 dilution), phospho-Chk1 (Ser345, 2348 S; Cell Signaling Technology; 1: 1000 dilution), phospho-RPA32 (S4/S8, A300-245A, Bethyl, 1: 1000 dilution) γ-tubulin (GTU-88; Sigma Aldrich; 1: 5000 dilution), GAPDH (0411, Santa Cruz; 1: 10000 dilution), LaminB1 (ab16048; Abcam; 1: 10000 dilution), A custom RHINO polyclonal antibody was outsourced from GenScript using the recombinant protein described in the “Protein purification section” with 6xHIS tag retained and produced in rabbit (GenScript; 1:1000 dilution). The secondary antibodies were mouse IgG HRP-linked (NA931, GE Healthcare; 1:5000) or rabbit IgG HRP-linked (NA934, GE Healthcare; 1:5000).

Antibodies used for immunofluorescence included phospho-H2AX (S139, 05-636 l; Millipore; 1: 5000 dilution), MYC (9B11; Cell Signaling; 1: 500 dilution), 53BP1 (100-304, Novus Biologicals; 1: 1000 dilution). Secondary antibodies were conjugated with Alexa Fluor 488 (Life Technologies), Alexa Fluor 568 (Life Technologies), or Cy5 (Life Technologies) and diluted 1:500.

### siRNA transfection

For DLD1 experiments, 125,000 cells were reverse transfected with 30 pmol of a pool of siRNAs (Dharmacon, SMARTpool, L-003462-00-0005) targeting *BRCA2* or a pool of scrambled (siNT, Dharmacon, SMARTpool, D-001810-10-05 or esiBRCA2 from Sigma, EHU031451) control sequences with RNAimax kit (Invitrogen), according to the manufacturer’s instructions. Halo-Polθ cells were nucleofected for live-cell mitotic imaging using a Lonza nucleofector with the indicated siRNAs targeting *BRCA2* and *RHNO1* using ∼200 ng siRNA/100,000 cells. To knock down *RHNO1*, U2OS cells were nucleofected with two separate siRNA (hs.Ri.RHNO1.13.1 and hs.Ri.RHNO1.13.2 from IDT).

### shRNA interference

shRNAs were cloned into a pLKO.1-Puro backbone as AgeI–EcoRI dsDNA oligomers (Integrated DNA Technologies) and introduced by four lentiviral transduction at 4-h intervals in the presence of 8 μg ml−1 polybrene (MilliporeSigma) using supernatant from transfected HEK293T cells. MEFs cells were selected with 3 μg/ml puromycin for 3 days and recovered for an additional day before evaluating the silencing efficiency. As a control, MEFs cells were infected with a pLKO.1-Puro plasmid encoding a scrambled shRNA sequence (Addgene plasmid #1864). A list of the shRNA sequences is provided in Supplementary Table 2.

### Lentiviral transduction

Lentiviral particles were produced in HEK293T cells in 10-cm plates by co-transfection of packaging vectors VSV-G, pMDLg/RRE, and pRSV-REV along with 10 μg of the desired plasmid using polyethylenimine (PEI). Virus-containing supernatant was collected ∼48-72 h post-transfection, cleared through a 0.2-μm filter, supplemented with 8 μg/mL polybrene (Sigma), and used for transduction. The medium was refreshed 6 to 12 h later.

### RT–qPCR

Total RNA was purified with the NucleoSpin RNA Clean-up (Macherey-Nagel) following the manufacturer’s instructions. Genomic DNA was eliminated by on-column digestion with DNase I. A total of 1 μg of RNA was reverse transcribed using iScript Reverse Transcription Supermix (Biorad), and cDNA was diluted 1:5 or 1:10. Reactions were run with ssoAdvanced SYBR Green Supermix (BioRad) with standard cycling conditions. Relative gene expression was normalized using *ACT1* or *GAPDH* as a housekeeping gene, and all calculations were performed using the ΔΔCt method.

### TLR assay

Lentiviral constructs coding for TLR (#31482) and I-SceI with donor e-GFP (#31476) were purchased from Addgene. To avoid the confounding effect of NHEJ on the repair of I-SceI-induced DNA breaks, we stably integrated the TLR construct into *LIG4*^*-/-*^ HEK293T. Cells were transduced with Cas9-RNP with sgRNAs against *TP53* and *9-1-1/RHNO1*, followed by virus particles of the I-SceI with donor e-GFP. Cells were collected 72 h later and analyzed on a BD LSRFortessa™ Cell Analyzer (BD Biosciences). Data were analyzed using FloJo software v.10 (TreeStar).

### CRISPR/Cas9-mediated gene knockout

To generate *LIG4* knockout, 10^6^ cells were transfected with 2.5 ugs of two plasmids containing Cas9 and the gRNA against exon4 in LIG4. Single clones were genotyped by PCR with primers amplifying a 600-bp sequence around the cutting sites to ascertain the presence of insertions or deletions (indels). Clones with possible homozygote status were amplified and subjected to confirmation by western blotting. *TP53*^*-/-*^ cells were generated by delivering the ribonucleoprotein complex of a single gRNA targeting exon 4 and Cas9 protein (IDT). Cells that survived a 7-day treatment of 10 μM Nutlin-3 were used for the Western blot for p53 expression. *RHINO1*^*-/-*^ cells were generated by delivering the ribonucleoprotein complex of a single gRNA targeting exon 3 and Cas9 protein (IDT). PCR genotyped single clones with primers amplifying a 600-bp around the cutting sites to ascertain the presence of insertions or deletions (indels). A list of the gRNA sequences is provided in Supplementary Table 2.

### Genome-wide CRISPR/Cas9 screens

CRISPR screens were performed as described(Hart *et al*., 2017). DLD1 cells were transduced with the lentiviral TKOv3 library at a low MOI (∼0.2–0.3) and selected with 4 ug/ml of puromycin for 48 h post-transduction. This was considered the initial time point (day 0). Cells with different genetic backgrounds were grown for additional 14 doublings. Cell pellets were frozen at each time point for genomic DNA (gDNA) isolation. A library coverage of ≥500 cells per sgRNA was maintained at every step. gDNA from cell pellets was isolated using the Quick-DNA Midiprep Plus Kit (ZymoResearch, D4075) and genome-integrated sgRNA sequences were amplified by PCR using the Q5 Mastermix (New England Biolabs Next UltraII). i5 and i7 multiplexing barcodes were added in a second round of PCR, and final gel-purified products were sequenced on Illumina HiSeq2500 or NextSeq500 systems to determine sgRNA representation in each sample. BAGEL2 was used to identify essential genes (Kim & Hart, 2021).

### Clonogenic and proliferation assays

500-1000 cells were seeded in 6-well plates for clonogenic assays. For IR sensitivity assays, cells were irradiated with 2 Gy collected after 30 min, counted, and seeded. After 10 days medium was removed, cells were rinsed with PBS and fixed with 4% paraformaldehyde for 10 minutes. Colonies were then rinsed with PBS and stained with 0.4% (w/v) crystal violet in 20% (v/v) methanol for 5 minutes. The stain was aspirated, and plates were rinsed in water three times and air-dried. Colonies were counted with the Fiji ImageJ plugin “Colony Area”. Data were plotted as surviving fractions relative to untreated cells. For the clonogenic assay on mitotic cells, cells were synchronized as described in Fig4A, and after irradiation, mitotic cells were recovered by mitotic shake-off.

For proliferation assay, an IncuCyte Live-Cell Analysis Imager (Essen/Sartorius) was employed to monitor confluency over time. Cell confluency was monitored every 24 h.

### IF-FISH

IF-FISH was performed as previously described (Sfeir & de Lange, 2012). Briefly, MEFs grown on coverslips were fixed with 4% paraformaldehyde for 10 min, washed with PBS, and permeabilized with 0.5% Triton X-100 buffer (0.5% Triton X-100, 20mM HEPES pH 7.9, 50mM NaCl, 3mM MgCl_2_, 300mM sucrose. Cells were then incubated in blocking solution (1mg/mL BSA, 3% goat serum, 0.1% Triton X-100, 1mM EDTA in PBS) at room temperature for 30 min, followed by incubation with primary antibody diluted in blocking solution for 2 hours at room temperature. After washing with PBST (0.1% Tween 20 in PBS) 3 times for 5 minutes each, cells were incubated with Alexa Fluor labeled secondary antibody (Thermo Fisher Scientific) for 45 minutes at room temperature. After washing with PBST 3 times for 5 minutes each, cells were dehydrated with ethanol series (70%, 95%, then 100%) and hybridized with TAMRA-OO-(TTAGGG)3 PNA probe (Applied Biosystems) in formamide hybridization solution (70% formamide, 0.5% blocking reagent (Roche), 10 mM Tris-HCl, pH 7.2) at 80°C for 5 min. Cells were allowed to cool at room temperature for 2 hours, then washed 4 times for 10 minutes each with formamide washing solution (70% formamide, 10mM Tris-HCl pH 7.2). Cells were washed with PBS 3 times for 5 min each, counterstained with DAPI, and coverslips were mounted on slides with anti-fade reagent (Prolong Gold, Invitrogen). Images were captured using a Nikon Eclipse 55i upright fluorescence microscope at 60–100× magnification and analyzed with Nikon software.

### Immunofluorescence

For the indicated treatments, cells were plated on 12-mm circular glass coverslips (Fisher Scientific), and immunofluorescence was performed using standard techniques. In brief, cells were fixed with 4% paraformaldehyde for 15 min at room temperature. Cells were washed with PBS, permeabilized with 0.5% (v/v) Triton X-100 for 10 min, and blocked for 30 min with PBS containing 3% goat serum (MilliporeSigma), 1 mg/ml bovine serum albumin (BSA, MilliporeSigma), 0.1% Triton X-100 and 1 mM EDTA. Cells were incubated with the same buffer containing primary antibodies for 1 hour at room temperature, followed by incubation with secondary antibodies overnight at 4C. DNA was counterstained with 5 μg/ml DAPI. Cells were mounted with ProLong Gold Antifade (Thermo Fisher Scientific), imaged on a Nikon Ti2 Eclipse upright fluorescence microscope at 40× magnification, and analyzed using Fiji ImageJ software. We employed a Nikon CSU-W1 Spinning Disk Confocal microscope with a 100x oil lens to image RHINO foci. Quantification of γH2AX in irradiation experiments was done using a script written in Fiji, briefly defining the ROI with the pH3S10 staining and quantifying the total integrated density of γH2AX staining (the sum of the values of the pixels in the image). The quantification of γH2AX foci in Olaparib-treated cells or RHINO foci in mitosis was done manually with blinded investigators.

### Western blotting analysis

Cells were collected by trypsinization and lysed in RIPA buffer (25 mM Tris-HCl pH 7.6, 150 mM NaCl, 0.1% SDS, 1% NP-40, 1% sodium deoxycholate). After two cycles of water-bath sonication at medium settings, lysates were incubated at 4°C on a rotator for an additional 30 min. Lysates were clarified by centrifugation for 30 min at 14,800 rpm at 4°C, and the supernatant was quantified using the enhanced BCA protocol (Thermo Fisher Scientific, Pierce). Equivalent amounts of proteins were separated by SDS–PAGE and transferred to a nitrocellulose membrane. Membranes were blocked in 5% milk in TBST or 5% BSA in TBST in the case of phosphorylated proteins for at least one hour at room temperature. Incubation with primary antibodies was performed overnight at 4°C. Membranes were washed and incubated with HRP-conjugated secondary antibodies at 1:5000 dilution, developed with Clarity ECL (Biorad), and acquired with a ChemiDoc MP apparatus (Biorad) and ImageLab v.5.2. Antibodies against LaminB1, GAPDH, or γ-tubulin were used as loading controls. We prepared SDS-PAGE gel using 25 μM of Phosbind Acrylamide (APExBIO) to resolve phosphorylated protein on a Western blot and 100 μM of MnCl_2_. To detect Halo-Polθ, cells were labeled with 150 nM JFX650-HaloTag ligand for 30 minutes, harvested, lysed in 2x Laemmli buffer, and loaded onto a 4-15% Mini-PROTEAN TGX Stain-Free polyacrylamide gel. Fluorescently labeled protein was detected on a BioRad Chemidoc using the Cy5.5 filter. Protein loading was detected using the Stain-Free filter on a BioRad Chemidoc after 45-second UV activation.

### Protein expression and purification

To test whether RHINO and Polθ directly interact, we outsourced purified recombinant human RHINO protein from GenScript. Rhno1 cDNA with N-terminal 6xHIS tag and TEV cleavage sequence was cloned into a pET30a vector and expressed in *E. coli* Artic Express (DE3). The cell lysate supernatant was isolated by ultracentrifugation and then applied to a Ni(II) chelate column. The target protein was further purified using a Superdex 200 column. The resulting purified protein was subjected to TEV protease cleavage to remove the 6xHIS tag and cleared via Ni(II) chelate column for *in vitro* assays. Purified Polθ was outsourced from Repare Therapeutics and purified as previously described (Monica Bubenik*, 2022). Purified 9-1-1 was kindly provided by the laboratory of Dr. Dirk Remus and purified as previously described (Castaneda *et al*, 2022).

### *In vitro* immunoprecipitation

Dynabeads Protein A was resuspended in a solution of 5ug RHINO antibody:200ul PBS 0.1% tween (PBST) per 50ul of beads and incubated with rotation at room temperature for 1 hour. The supernatant was removed, and conjugated beads were washed with PBST with gentle pipetting. Beads were resuspended with PBST to make a 50% slurry. Before immunoprecipitation, 10ul of packed beads were washed once with the reaction buffer (25mM HEPES KOH pH 7.6, 150mM NaCl, 0.02% NP40, 1mM EDTA, 5% glycerol, 0.1mg/mL BSA, 1mM DTT). One μM purified RHINO, 1 μM purified 9-1-1, and 0.311uM purified Polθ were used for in vitro immunoprecipitation experiments. 1X Master Mix was added to the conjugated beads and the appropriate amount of purified protein to a total volume of 100ul. The immunoprecipitation reaction was carried out at 37C for 30 minutes with gentle agitation. The supernatant was cleared, and the beads were washed twice with 1X Master Mix. Immunoprecipitated proteins were eluted from the beads using 50mM glycine pH 2.8 for 20 minutes at room temperature with gentle agitation and immediately neutralized with Tris pH 8.0. Laemmli buffer was added to 25% of the eluant, boiled at 95C for 5 minutes, loaded into SDS-PAGE gels, and subjected to western blot.

### Coomassie stain

Purified Polθ, 9-1-1, and RHINO proteins were diluted to the concentrations reflecting those in the in vitro immunoprecipitation assay and boiled with Laemmli buffer for 5 minutes at 95C. Proteins were loaded in a 4-20% gradient SDS-PAGE gel and subjected to Coomassie staining. The SDS-PAGE gel was fixed for 1 hour in 50% ethanol:10% acetic acid solution and then washed overnight in 50% methanol:10% acetic acid. After, the gel was stained in Coomassie stain (0.1% w/v Coomassie blue R250, 50% methanol:10% acetic acid) for 4 hours at room temperature. The gel was washed until the stain was removed using 50% methanol:10% acetic acid. Then, the gel was incubated with a 5% acetic acid solution for 1 hour before imaging. All were performed at room temperature with gentle agitation.

### Coimmunoprecipitation (CoIP)

HEK293T cells were co-transfected with the indicated plasmid constructs using polyethyleneimine (PEI). 48hr post-transfection cells were harvested in cold PBS, washed once with PBS, and lysed for 15min on ice in 500 μL Lysis Buffer (50mM Tris-HCl pH 7.4, 150mM NaCl, 1% Triton X-100, 0.05% SDS, 1mM EDTA, 1mM DTT, 1mM PMSF, and protease inhibitor cocktail). Salt boost was performed by adding 25 μL 5M NaCl and incubating on ice for an additional 5min. Then, 500 μL cold H_2_O was added, and lysates were spun at max speed for 10min to pellet debris. 60ul of protein G Sepharose beads conjugated to M2 Flag antibody (Millipore Sigma Cat. M8823-5ML) was added to each sample. Samples were incubated for 3 hours, rotating at 4°C. Beads were washed 4 times in Wash Buffer (50mM Tris-HCl pH 7.4, 150mM NaCl, 1% Triton X-100, 0.1% SDS), and proteins were eluted in 2X Laemmli buffer by boiling at 95°C for 5min for Western blot.

### Immunoprecipitation

DLD1 RHINO-MYC-FLAG cells were harvested by either trypsinization or mitotic shake-off, washed once with PBS, and resuspended in 1mL of IP lysis buffer (50mM Tris-HCl pH 7.5, 150mM NaCl, 0.5% NP-40, 0.1% sodium deoxycholate, 5% glycerol, 1mM EDTA), supplemented with 1X complete protease inhibitor cocktail, 1mM DTT, and 375U/mL Benzonase nuclease (Sigma, E1014). Cells were incubated on ice for 1 hour, and lysates were cleared by centrifugation at max speed at 4C for 15 min. 3 mg of cell extract was incubated with 60 μl/sample of Anti-FLAG® M2 Magnetic Beads (Millipore Sigma, M8823-1ML) overnight on a rotating wheel at 4C. Immunoglobulin-antigen complexes were washed 3 times for 15 min in cold TBS before elution (1X TE, 1% SDS).

### Cell cycle analysis by FACS

0.5 × 10^6^ cells were collected by trypsinization, washed with cold PBS, and fixed with 70% ice-cold ethanol at 4°C for at least 24h. After fixation, cells were washed once with PBS, and DNA was labeled with propidium iodide (50 μg/ml) in the presence of RNase A (0.2 mg/ml, Qiagen) and Triton X-100 (0.01%) for 30 minutes at room temperature. Data on DNA content were acquired on a BD LSRFortessa™ Cell Analyzer (BD Biosciences) using BD FACSDiva™ Software (BD Biosciences) and analyzed using FlowJo v.10 (TreeStar).

### Live cell imaging microscopy

Cells were plated onto glass-bottom 24-well plates for live-cell imaging experiments and imaged with an Olympus IX83 inverted microscope. Imaging was performed at 37C and 5% CO2 with the 100x objective and the 640 nm laser with highly inclined laminated optical sheet illumination (lightly angled 0.5). Thymidine block was performed using 2 mM Thymidine and 9 μM RO-3306 was used to synchronize cells at the G2/M boundary. Cells were treated with 1 μM olaparib for ∼16 hours or 20 μg/mL Zeocin for one hour. For mitotic imaging, cells were released in 100 ng/mL nocodazole media. Halo-Polθ cells were labeled with JFX650 at 150 nM for 30 – 60 minutes. After labeling, cells were washed three times with complete medium and allowed to rest for 10 minutes before adding full fresh medium. Z-stack images were acquired at 150 nm intervals. Mitotic Polθ foci were counted by visual inspection in ImageJ. They were classified as foci if a particle remained static and could be visualized through at least four consecutive Z-frames.

### Statistical analysis

All statistical analysis was performed with GraphPad Prism 9. Sample sizes and the statistical tests used are specified in the figure legends.

