## Supplemental Figures for "RHINO restricts MMEJ activity to mitosis"

### Figure S1

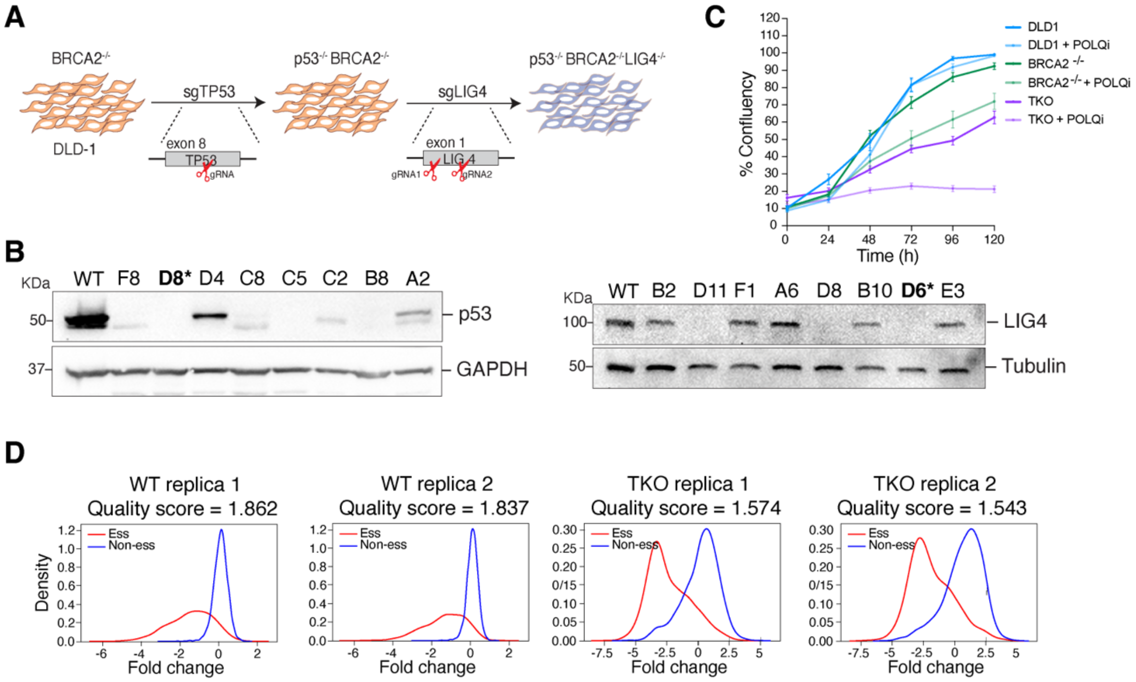


**E**

BF score (genes SL with BRCA2)

| | WT | TKO |
|---|---|---|
| APEX2 | -16.3 | 23.6 |
| FEN1 | 2.0 | 12.9 |
| ALC1 | -29 | 15.5 |
| CIP2A | -23.4 | 18.3 |
| RNASEH2A | -6.9 | 26.5 |
| RNASEH2B | -7.38 | 13.5 |
| RNASEH2C | 1.3 | 9.6 |

**F**

BF score (known MMEJ factors)

| | WT | TKO |
|---|---|---|
| POLQ | 8.7 | 20.9 |
| PARP1 | -26.6 | 20.5 |
| HMCES | -13.9 | 7.3 |

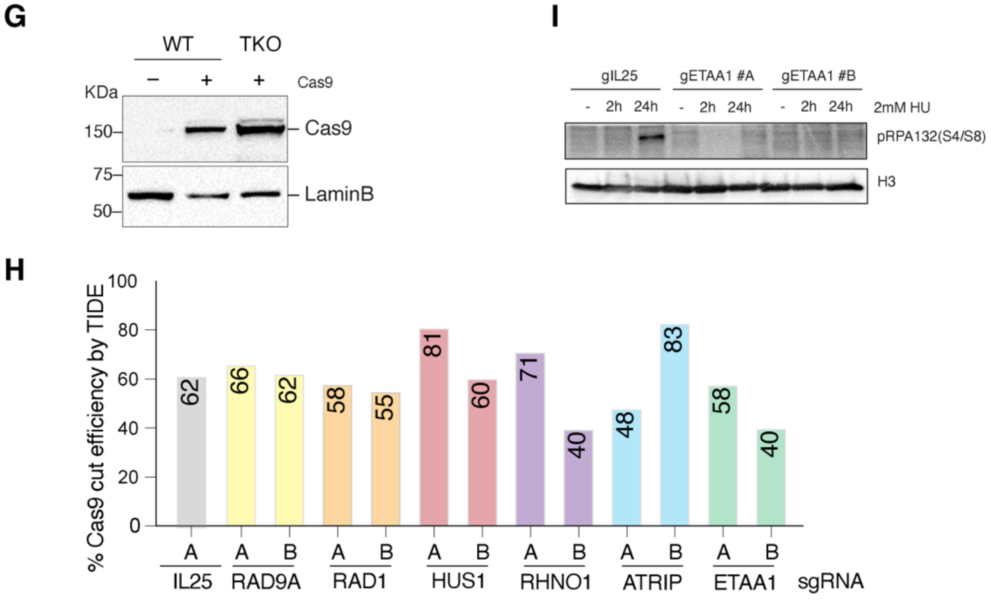

**Fig. S1. The 9-1-1/RHINO complex is a novel MMEJ factor.**

(**A**) Schematic of CRISPR-Cas9 strategy for sequential knock out of *TP53* and *LIG4* from DLD1 cells. (**B)** Western blot of DLD1 cells transduced with sgRNA against *TP53* or *LIG4*. Lysates were probed with antibodies detecting p53 and GAPDH (loading control) or against Lig4 and Tubulin (loading control). (**C**) Growth curve using IncucyteS3 in the indicated cells treated with Polθ inhibitor RP6685. (**D**) Kernel density estimates reference core-essential genes (red) and non-essential genes (blue) in each CRISPR/Cas9 screen replicate. The quality score was calculated using Cohen's D statistic, which represents the mean log fold change difference between core-essential and non-essential genes divided by the pooled standard deviation of the log fold change of all reference genes(*28*). (**E**) BF scores for selected genes known to be essential for the survival of *BRCA2*$^{-/-}$ cells. SL = synthetic lethal. (**F**) BF scores for known MMEJ genes*.* (**G**) Western blot analysis to monitor Cas9 levels in WT and TKO cells. Lamin B1 antibody is used as a loading control. (**H**) Editing efficiency of selected sgRNAs. Genomic DNA was isolated from WT cells expressing Flag-Cas9 and the indicated sgRNAs. The region flanking the sgRNA cut site was amplified by PCR and sequenced. Results from the TIDE analysis (*62*) are shown. (**I**) Functional validation of *ETAA1* depletion. Cells expressing Flag-Cas9 and the indicated sgRNAs were treated for 2 and 24 hours with 2 mM hydroxyurea (HU). Western blot analysis was performed for pRPA (S4/S8). H3 was used as a loading control.

Figure S2

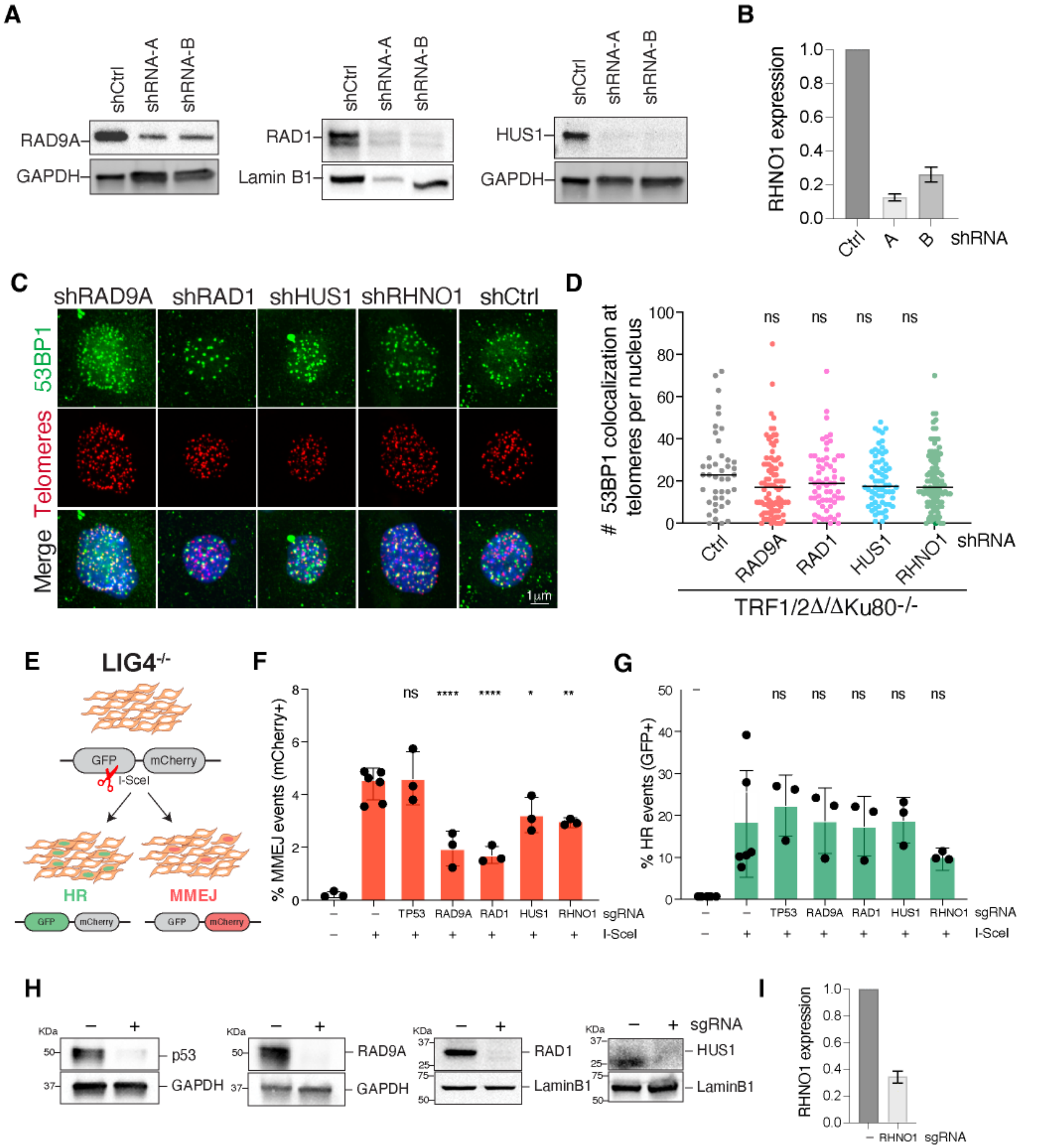


**Fig. S2. The role of 9-1-1/RHINO in MMEJ is ATR-signaling independent.**

(**A**) Western blot for 9-1-1 complex members in *TRF1/2*$^{\Delta/\Delta}$*Ku80*$^{-/-}$*p53*$^{-/-}$ MEFs after shRNA knockdown. GAPDH or Lamin B1 antibodies are used as a loading control. (**B**) qPCR analysis of *RHNO1* mRNA expression after shRNA knockdown. Relative gene expression was normalized using TBP as a housekeeping gene. Data are the mean of two independent experiments. (**C**) Representative images of IF-FISH analysis of telomere dysfunction-induced foci (TIFs) in *TRF1/2*$^{\Delta/\Delta}$*Ku80*$^{-/-}$*p53*$^{-/-}$ cells depleted for 9-1-1 and RHNO1. Telomeres are stained with a FISH probe (red), and 53BP1 is stained using an antibody (green). DNA is stained with DAPI (blue). (**D**) DNA damage was assessed by quantifying the colocalization of 53BP1 with telomeres (TTAGGG) in *TRF1/2*$^{\Delta/\Delta}$*Ku80*$^{-/-}$*p53*$^{-/-}$ cells with the indicated treatment. Each dot on the graph represents a single cell. The graph represents at least 40 nuclei from 2 independent experiments, and black lines indicate the mean. Statistical analysis was performed using one-way ANOVA. (**E**) Diagram of the Traffic Light Reporter (TLR) in NHEJ deficient cells (*LIG4*$^{-/-}$). The schematic represents the different outcomes after an I-SceI-induced DSB. When a break is resolved through HR, full-length eGFP will be reconstituted (GFP+). If the break is repaired via MMEJ, eGFP will be translated out of frame while mCherry sequences become in frame leading to red fluorescence (mCherry +). (**F**) Quantifying MMEJ events by TLR after depletion of 9-1-1 and RHINO through Cas9-RNP nucleofection. Statistical analysis was performed using one-way ANOVA (****$p<0.0001$, **$p<0.01$, *$p<0.05$). (**G**) Quantifying HR events by TLR (*35*) upon loss of 9-1-1 and RHINO. Statistical analysis was performed using one-way ANOVA. (**H**) Western blot for 9-1-1 complex subunits in HEK293T upon knockdown. GAPDH and Lamin B1 antibodies served as loading controls. (**I**) qPCR analysis of *RHNO1* mRNA expression after knockdown in cells used for the TLR assay. Relative gene expression was normalized using *ACT1* as a housekeeping gene.

### Figure S3

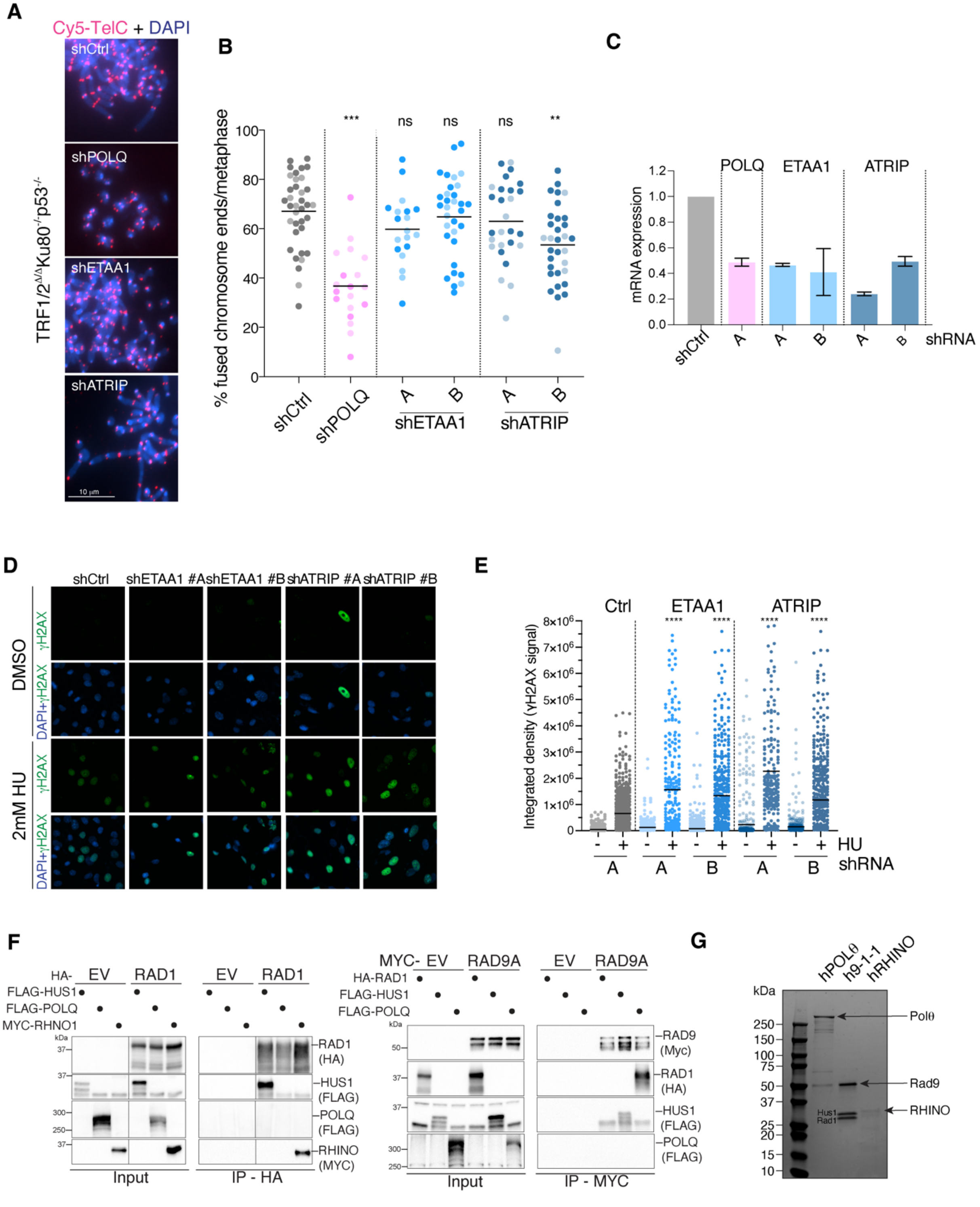
A
Cy5-TelC + DAPI
shCtrl
shPOLQ
shETAA1
shATRIP
TRF1/2^Δ/Δ Ku80^-/- p53^-/-
10 μm
B
% fused chromosome ends/metaphase
*** ns ns ns **
shCtrl
shPOLQ
A B
shETAA1
A B
shATRIP
C
POLQ ETAA1 ATRIP
mRNA expression
shCtrl A A B A B shRNA
D
shCtrl shETAA1 #A shETAA1 #B shATRIP #A shATRIP #B
DMSO
2mM HU
γH2AX
DAPI+γH2AX
E
Ctrl ETAA1 ATRIP
Integrated density (γH2AX signal)
**** **** **** ****
HU
A A B A B shRNA
F
HA- EV RAD1
FLAG-HUS1
FLAG-POLQ
MYC-RHNO1
kDa
RAD1 (HA)
HUS1 (FLAG)
POLQ (FLAG)
RHINO (MYC)
Input
IP - HA
MYC- EV RAD9A
HA-RAD1
RAD9 (Myc)
IP - MYC
G
kDa
hPOLθ
h9-1-1
hRHINO
Polθ
Rad9
Hus1
Rad1
RHINO

**Fig. S3. 9-1-1/RHINO function in ATR signaling is insufficient to promote MMEJ.**

(**A**) Representative images of a metaphase spread from *TRF1/2*$^{\Delta/\Delta}$*Ku80*$^{-/-}$*p53*$^{-/-}$ cells depleted for *ETAA1*, *ATRIP* or *POLQ*. Telomeres are marked by FISH using a Cy5-[CCCTAA]3 PNA probe (red), and chromosomes are counterstained with DAPI (blue). White arrows indicate examples of telomeric fusions in the control sample. (**B**) Quantification of telomere fusions mediated by MMEJ related to panel A. Data are the mean of two independent experiments. The different colors used for the dots represent the results from independent experiments. (**C**) qPCR analysis of RHNO1 mRNA expression after knockdown with shRNA in *TRF1/2*$^{\Delta/\Delta}$ *Ku80*$^{-/-}$*p53*$^{-/-}$ cells used for the metaphase spreads. Relative gene expression was normalized using ACT1 as a housekeeping gene. (**D**) Functional validation of ETAA1 and ATRIP depletion in *TRF1/2*$^{\Delta/\Delta}$*Ku80*$^{-/-}$*p53*$^{-/-}$ cells. Cells with the indicated shRNAs were treated for 3 hours with 2mM hydroxyurea (HU) and subjected to IF analysis for γH2AX staining. Representative images from one of two independent experiments. (**E**) Quantification of γH2AX signal in two independent experiments. Data are the mean of two independent experiments and statistical analysis performed using one-way ANOVA between the HU-treated samples. (**F**) Whole-cell extracts from HEK293T cells co-transfected with plasmids expressing FLAG-POLQ and subunits of 9-1-1 (MYC-RAD9, HA-RAD1, and FLAG-HUS1) were subjected to immunoprecipitation followed by western blot with the indicated antibodies. (G) Representative Coomassie-stained SDS-PAGE analysis of purified human Polθ, 9-1-1 complex, and RHINO.

### Figure S4

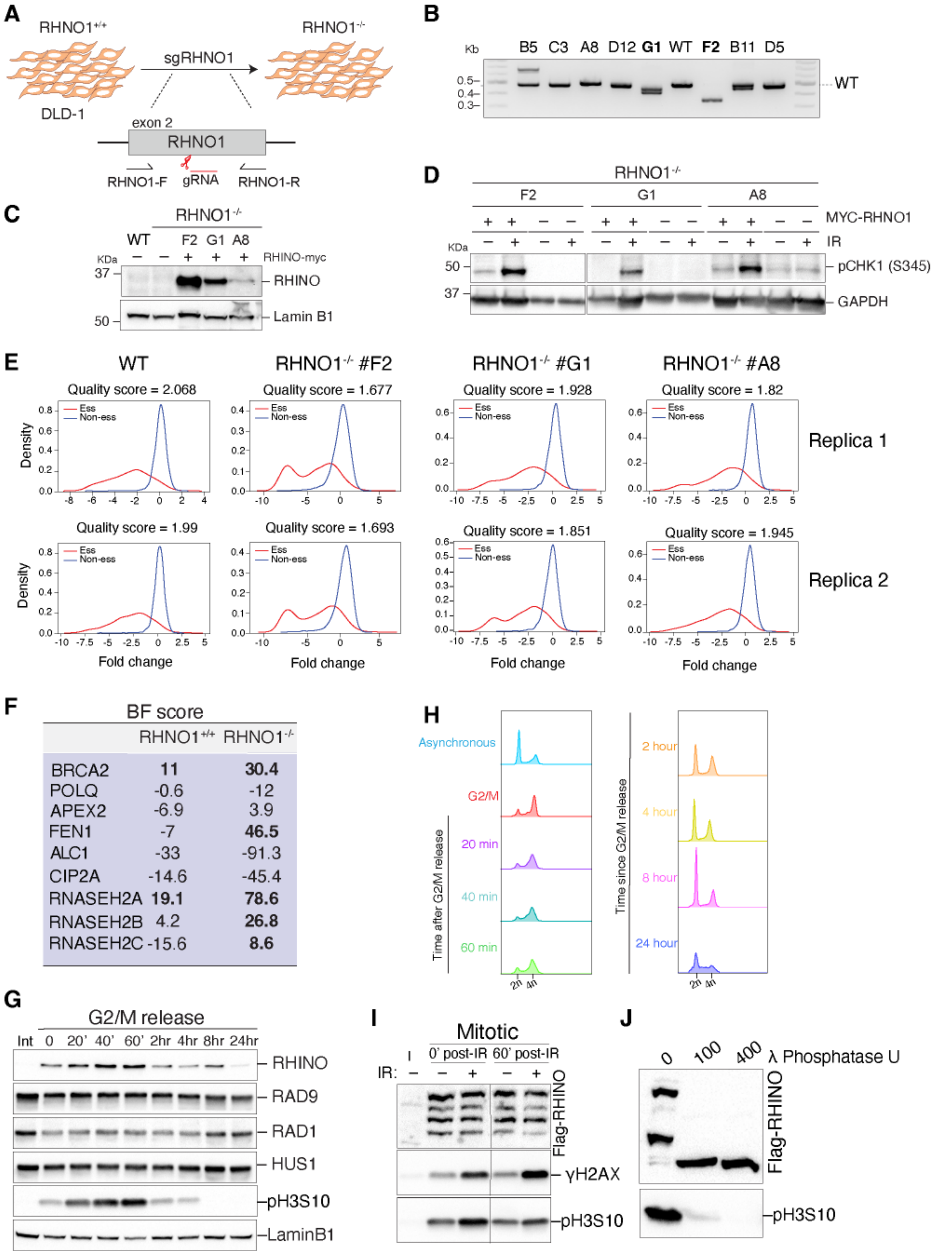
A
RHNO1+/+
sgRHNO1
RHNO1-/-
DLD-1
exon 2
RHNO1
RHNO1-F
gRNA
RHNO1-R
B
Kb
B5 C3 A8 D12 G1 WT F2 B11 D5
0.5
0.4
0.3
WT
C
RHNO1-/-
WT F2 G1 A8
KDa
RHINO-myc
37
RHINO
50
Lamin B1
D
RHNO1-/-
F2 G1 A8
MYC-RHNO1
IR
KDa
50
pCHK1 (S345)
37
GAPDH
E
WT
RHNO1-/- #F2
RHNO1-/- #G1
RHNO1-/- #A8
Quality score = 2.068
Quality score = 1.677
Quality score = 1.928
Quality score = 1.82
Replica 1
Quality score = 1.99
Quality score = 1.693
Quality score = 1.851
Quality score = 1.945
Replica 2
Ess
Non-ess
Density
Fold change
F
BF score
RHNO1+/+ RHNO1-/-
BRCA2 11 30.4
POLQ -0.6 -12
APEX2 -6.9 3.9
FEN1 -7 46.5
ALC1 -33 -91.3
CIP2A -14.6 -45.4
RNASEH2A 19.1 78.6
RNASEH2B 4.2 26.8
RNASEH2C -15.6 8.6
H
Asynchronous
G2/M
20 min
40 min
60 min
Time after G2/M release
2n 4n
2 hour
4 hour
8 hour
24 hour
Time since G2/M release
G
G2/M release
Int 0 20' 40' 60' 2hr 4hr 8hr 24hr
RHINO
RAD9
RAD1
HUS1
pH3S10
LaminB1
I
Mitotic
0' post-IR 60' post-IR
IR:
Flag-RHINO
γH2AX
pH3S10
J
0 100 400 λ Phosphatase U
Flag-RHINO
pH3S10

**Fig. S4. RHINO is strictly expressed in mitosis.**

(**A**) Schematic of the CRISPR/Cas9 strategy to knockout RHNO1. (**B**) Genotyping PCR on genomic DNA extracted from cells targeted with CRISPR/Cas9 and detecting deletions in the *RHNO1* gene. Clones employed on the screen are highlighted in bold. (**C**) Western blot of RHNO1$^{-/-}$ clones complemented with RHINO-Myc-Flag expressing vector. Lamin B1 is used as a loading control. (**D**) Functional validation of RHINO deletion in three clones used in the CRISPR/Cas9 screen based on the accumulation of phospo-CHK1 following irradiation. Three RHNO1$^{-/-}$ clones were complemented with RHINO-MYC-FLAG expressing vector, subjected to western blot 3 hours post-irradiation (2 Gy), and monitored for phospo-CHK1. GAPDH antibody is used as a loading control. (**E**) Kernel density estimates of reference core-essential genes (red) and non-essential genes (blue) in each replica of the DLD1 CRISPR/Ca9 screen. (**F**) BF scores for selected genes known to be synthetic lethal with BRCA2 null cells. (**G**) Western blot analysis of 9-1-1/RHINO expression during the cell cycle. Cells overexpressing RHINO-MYC-FLAG underwent a double thymidine block and were then released in CDK1 inhibitor for 16h to induce a G2/M arrest. Cells were collected at the indicated time points and subjected to western blot to detect RHINO and components of the 9-1-1 complex. The phosphoantibody against serine 10 in H3 (pS10H3) was used as a mitotic marker. Lamin B1 was used as a loading control. Int = interphase. (**H**) Cell cycle analysis by FACS of the samples analyzed in panel G. (**I**) Phosho-tag gel of DLD1 cells overexpressing RHINO-MYC-FLAG synchronized in mitosis and irradiated with 2 Gy. Cells were collected immediately after irradiation and after 60 minutes. γH2AX is used as a marker of DNA damage, and pS10H3 marks mitosis. (**J**) Phosho-tag gel of DLD1 cells expressing RHINO-MYC-FLAG synchronized in mitosis and treated with either 100 or 400 units of lambda ($\lambda$) phosphatase. pS10H3 is used as a marker of mitosis.

Figure S5

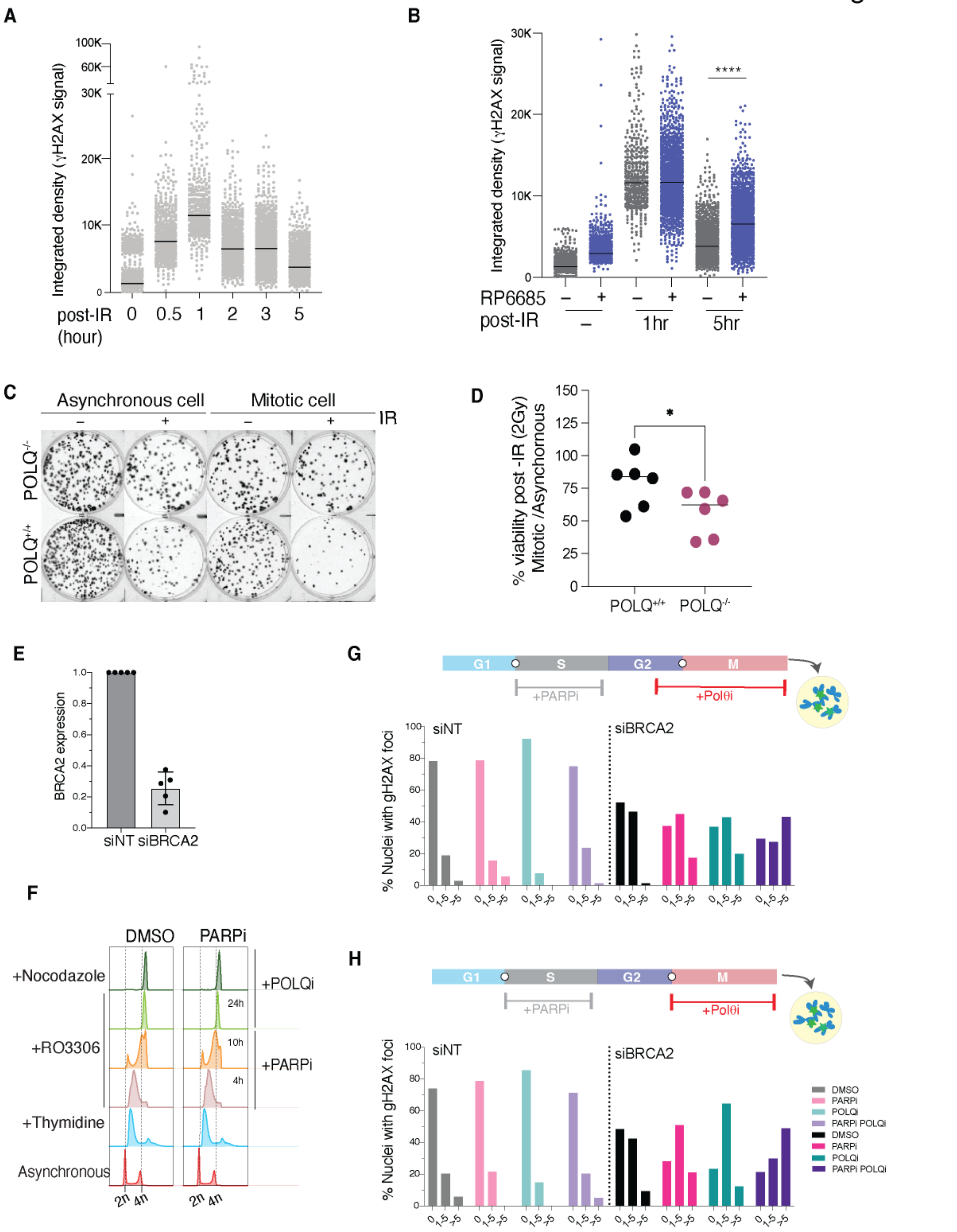
A
Integrated density (γH2AX signal)
post-IR (hour)
0 0.5 1 2 3 5
B
Integrated density (γH2AX signal)
RP6685
post-IR
1hr
5hr
****
C
Asynchronous cell
Mitotic cell
IR
POLQ-/-
POLQ+/+
D
% viability post -IR (2Gy) Mitotic /Asynchornous
POLQ+/+ POLQ-/-
E
BRCA2 expression
siNT siBRCA2
F
DMSO
PARPi
+Nocodazole
+RO3306
+Thymidine
Asynchronous
+POLQi
+PARPi
24h
10h
4h
2n 4n
G
G1 S G2 M
+PARPi
+Polθi
% Nuclei with gH2AX foci
siNT
siBRCA2
H
DMSO
PARPi
POLQi
PARPi POLQi

**Fig. S5. Polθ is the major mitotic DSB repair enzyme.**

(**A**) Quantification of γH2AX intensity signal from IF in control cells synchronized and irradiated with 2Gy in mitosis. IF were performed at the indicated time points. 0h is the no-irradiation control sample. At least 400 nuclei per sample were analyzed. (**B**) Quantification of γH2AX intensity signal by immunofluorescence in cells synchronized as in panel a but treated with Polθ inhibitor RP6685. (**C**) Representative images of clonogenic survival of *POLQ*$^{+/+}$ and *POLQ*$^{-/-}$ cells irradiated with 2Gy in asynchronous or mitotic cells. (**D**) The viability post-irradiation of mitotic cells was normalized to the asynchronous population. Bars represent the mean of 5 independent experiments. An unpaired t-test was run for statistical significance (*$p<0.05$). (**E**) qPCR analysis of *BRCA2* mRNA expression after siRNA knockdown of the samples analyzed in main figure 4-F and S5G-H. Relative gene expression was normalized using *ACT1* as a housekeeping gene. (**F**) Cell cycle analysis by FACS of the samples analyzed in Figure 4E. (**G-H**) Cells were synchronized at the G1/S interphase using a thymidine block and released into S-phase with a PARP inhibitor (Olaparib). PARP inhibitor was withdrawn upon exit from S phase, and CDK1 inhibitor was added (RO3306). Cells were then washed and treated with nocodazole, fixed, and stained with γH2AX antibody one hour after release into mitosis. Polθ inhibitor was added at the end of G2 and into M in panel (**G**) or exclusively in M (**H**).

### Figure S6

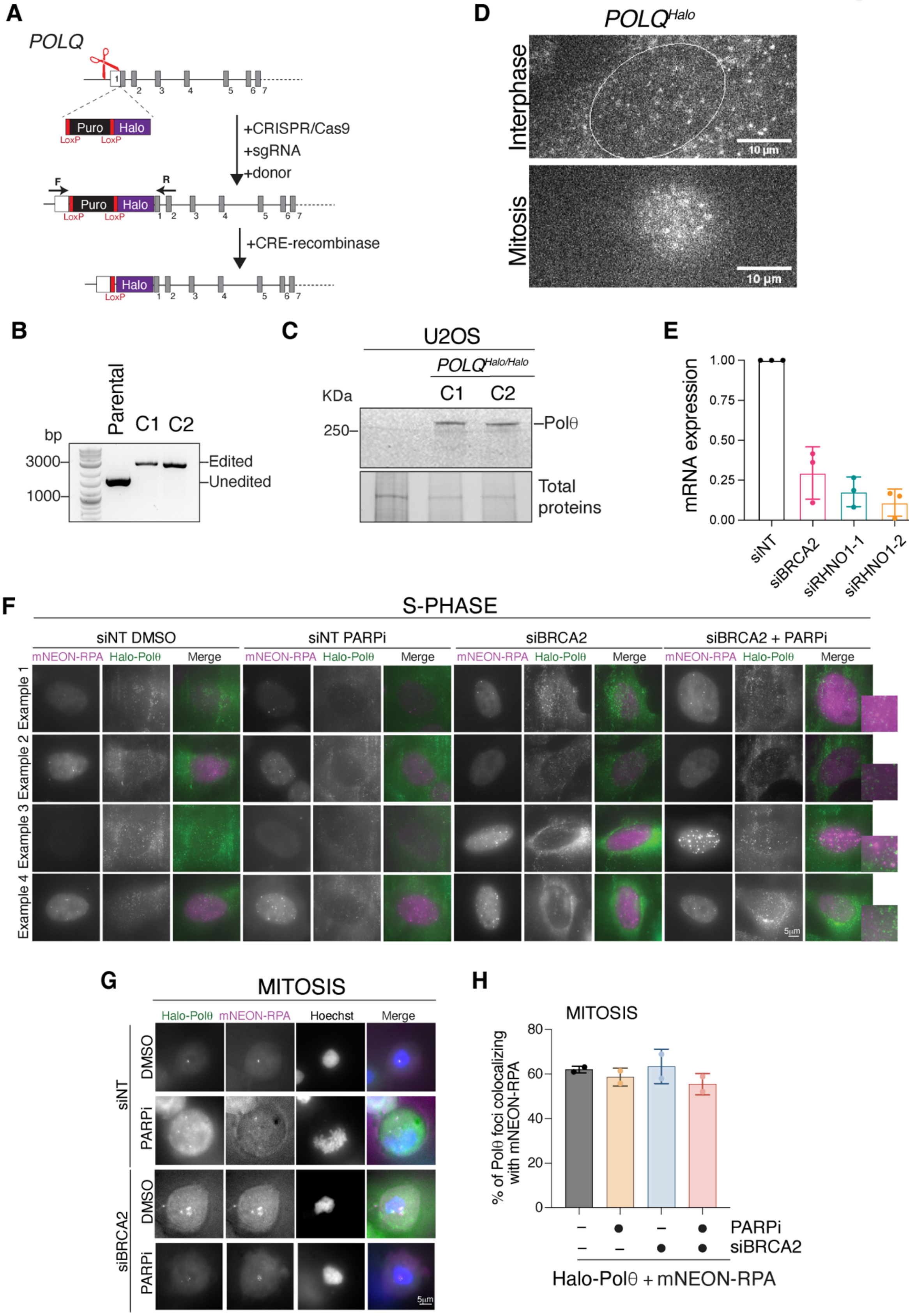
A
POLQ
Puro
Halo
LoxP
+CRISPR/Cas9
+sgRNA
+donor
F
R
+CRE-recombinase
D
POLQ^Halo
Interphase
Mitosis
10 µm
B
bp
Parental
C1
C2
3000
1000
Edited
Unedited
C
U2OS
POLQ^Halo/Halo
KDa
250
Polθ
Total proteins
E
mRNA expression
1.00
0.75
0.50
0.25
0.00
siNT
siBRCA2
siRHNO1-1
siRHNO1-2
F
S-PHASE
siNT DMSO
siNT PARPi
siBRCA2
siBRCA2 + PARPi
mNEON-RPA
Halo-Polθ
Merge
Example 1
Example 2
Example 3
Example 4
5µm
G
MITOSIS
Hoechst
siNT
siBRCA2
DMSO
PARPi
H
% of Polθ foci colocalizing with mNEON-RPA
80
60
40
20
0
PARPi
siBRCA2
Halo-Polθ + mNEON-RPA

**Fig. S6. RHINO recruits Polθ to DSB sites in mitosis.**

(**A**) Strategy for Halo-Polθ targeting using CRISPR/Cas9. (**B**) Genotyping PCR on genomic DNA extracted from targeted cells as in (A). Genomic PCR was conducted on cells upon excision of the Puromycin cassette with Cre. (**C**) Western blot analysis on two Halo-Polθ clones and the parental line as a negative control. Protein loading was detected using the Stain-Free filter on a BioRad Chemidoc. (**D**) Representative still image of Halo-Polθ in live-cell movies of interphase and mitotic cells. (**E**) qPCR analysis of *BRCA2* and *RHNO1* mRNA expression after siRNA knockdown of the samples analyzed in Fig. 5A-C. Relative gene expression was normalized using *GAPDH* as a housekeeping gene. (**F**) Representative images of Halo-Polθ and mNEON-RPA in live-cell experiment in S-phase cells. Cells treated with siRNA against control and *BRCA2* were transfected with a vector for mNEON-RPA32 and synchronized in G1/S using thymidine block and released into S-phase in the presence of PARP inhibitor (Olaparib, 10 μm). Images were taken 4 hours after release from the thymidine block. mNEON-RPA is depicted in magenta and Halo-Polθ in green. (**G**) Representative images of Halo-Polθ and mNEON-RPA in live-cell experiment in M-phase cells. Cells treated with siRNA against control and *BRCA2* were transfected with a vector for mNEON-RPA32 and synchronized in G1/S using thymidine block and released into S-phase in the presence of PARP inhibitor (Olaparib, 10 μm) followed by CDKi and release in nocodazole to enrich for mitotic cells. mNEON-RPA is depicted in magenta and Halo-Polθ in green. (**H**) Quantification of Halo-Polθ foci colocalization with mNEON-RPA foci. Bars represent the mean of 2 independent experiments. At least 20 nuclei were counted for each sample.
